# Novel mechanism of hypoxic neuronal injury mediated by non-excitatory amino acids and astroglial swelling

**DOI:** 10.1101/2022.05.10.491362

**Authors:** Iris Álvarez-Merz, Ioulia V. Fomitcheva, Jeremy Sword, Jesús M. Hernández-Guijo, José M. Solís, Sergei A. Kirov

**Author notes:** **Correspondence should be addressed to:** Sergei A. Kirov, PhD, **Email:**.

## Abstract

Bleeding into cerebral parenchyma during hemorrhagic stroke or head trauma leads to ischemia and release of plasmatic content, including amino acids (AA). Although excitotoxic AA have been extensively studied, little is known about non-excitatory AA during hypoxic injury. Hypoxia-induced synaptic depression becomes irreversible after adding non-excitatory AA to hippocampal slices, alongside their intracellular accumulation and increased tissue electrical resistance. A combination of four non-excitatory AA (L-alanine, glycine, L-glutamine, L-serine: AGQS) at plasmatic concentrations was applied to brain slices from transgenic mice expressing EGFP in pyramidal neurons or astrocytes during normoxia or hypoxia. Two-photon imaging, changes in light transmittance (LT), and electrophysiological field recordings followed by electron microscopy in hippocampal CA1 *st. radiatum* were used to monitor synaptic function concurrently with cellular swelling and injury. During normoxia, AGQS-induced increase in LT was due to astroglial but not neuronal swelling. Fast LT raise during hypoxia and AGQS manifested neuronal and astroglial swelling accompanied by a permanent loss of synaptic transmission and irreversible dendritic beading, signifying acute tissue damage. Neuronal injury was not triggered by spreading depolarization which did not occur in our experiments. Hypoxia without AGQS did not cause cell swelling, leaving dendrites intact. Inhibition of NMDA receptors prevented neuronal damage and irreversible loss of synaptic function. Deleterious effects of AGQS during hypoxia were prevented by alanine-serine-cysteine transporters (ASCT2) and volume-regulated anion channels (VRAC) blockers. Our findings suggest that swelling induced by intracellular accumulation of non-excitatory AA and release of excitotoxins through antiporters and VRAC may exacerbate the hypoxia-induced neuronal injury.

**Significance Statement:** Little is known if non-excitatory amino acids (AA) contribute to cellular injury when released during bleeding, as in hemorrhagic stroke and head trauma. Alanine, glycine, glutamine, and serine are one of the most abundant in plasma. Remarkably, during hypoxia, these non-excitatory AA caused severe neuronal and astroglial swelling and irreversible dendritic injury alongside a permanent loss of synaptic function. Activation of NMDA receptors was implicated in the onset of damage. Experimental evidence pointed to the involvement of alanine-serine-cysteine transporter 2 (ASCT2) and volume-regulated anion channels (VRAC) as molecular mechanisms underlying AA-induced damage during hypoxia. A detailed understanding of how brain injury evolves with non-excitatory AA during hypoxia will help design brain recovery treatments in neurological conditions involving bleeding.

## Introduction

The rupture of blood vessels during neurological emergencies such as hemorrhagic stroke and traumatic brain injury (TBI) leads to the release of plasmatic content, including amino acids (AA), to the cerebral parenchyma. Ischemia then quickly develops due to the reduced supply of oxygen and glucose to the affected cortical tissue. In these brain pathologies, 55-90% of patients have spreading depolarizations (SDs) that contribute to new lesion formation, penumbra expansion, and poor prognoses for patients’ recovery (Hartings et al., 2017). Still, even without SDs, hemorrhagic progression of contusions (HPC) after TBI propagates secondary brain cell damage hours and days after primary injury (Kurland et al., 2012; Adatia et al., 2021). Furthermore, in focal stroke, AA are released in the ischemic core from necrotic cells, contributing to penumbra expansion (Hossmann, 1994). Many mechanisms can contribute to secondary cell injury. The role of excitatory AA glutamate and aspartate that trigger excitotoxicity cascades with subsequent secondary neuronal damage has been extensively studied (Choi and Rothman, 1990; Yan et al., 2020). However, not much is known about the harmful effects of non-excitatory AA, especially when combined with hypoxia, in mediating the secondary injury cascades during these pathological conditions.

Extracellular and intracellular stores of AA are tightly controlled by diverse families of transporters (Bröer and Bröer, 2017). Pathologically high AA concentration in the cerebral parenchyma may lead to intracellular accumulation of AA through their transport systems, followed by osmotically obligated water and brain cell swelling. Cellular swelling can be indirectly measured by increased extracellular resistance (Chebabo et al., 1995) and field potential amplitude (Rosen and Andrew, 1990). Accordingly, we have shown that the intracellular accumulation of a mixture of seven non-excitatory AA (L-alanine, glycine, L-histidine, L-glutamine, L-serine, L-threonine, and taurine, abbreviated as one letter code: AGHQSTU) applied to hippocampal slices at their plasmatic concentrations induces synaptic potentiation suggesting cell swelling (Álvarez-Merz et al., 2021). Most of these AA are substrates of several Na^+^-dependent AA transporter systems located in the neuronal and astroglial plasma membrane, such as systems A and N (SLC38 family) (Bröer, 2014) and alanine-serine-cysteine transporters (ASCT; SLC1 family) (Bröer et al., 1999; Weiss et al., 2001; Gliddon et al., 2009). However, pharmacological inhibition of system A transport did not prevent synaptic potentiation induced by the AGHQSTU mixture (Álvarez-Merz et al., 2021), indicating that system A alone is insufficient for intracellular accumulation of AA observed in this study.

Volume-regulatory mechanisms are activated in response to changes in cell volume. Cell swelling opens volume-regulated anion channels (VRAC), leading to the efflux of inorganic and organic osmolites, including chloride and taurine, as well as glutamate, aspartate, and D-serine (Jackson and Strange, 1993; Hyzinski-García et al., 2014). The release of excitatory AA by activated VRAC was proposed to aggravate excitotoxicity and induce neuronal damage in metabolically compromised tissue (Wilson et al., 2019). Indeed, we have shown that the reversible loss of synaptic transmission induced by hypoxia turns irreversible by non-excitatory AA and involves NMDA receptor (NMDAR) activation (Álvarez-Merz et al., 2021).

Brain slices from transgenic mice with fluorescent neurons and astrocytes (Feng et al., 2000; Nolte et al., 2001) allow real-time imaging of cellular volume changes with 2-photon laser scanning microscopy (2PLSM) (Risher et al., 2009). Here, we used 2PLSM and electron microscopy (EM) to search for direct evidence of cellular injury during hypoxia by four non-excitatory AA (L-alanine, glycine, L-glutamine, and L-serine, abbreviated as AGQS) that are one of the most abundant AA in blood plasma (Lerma et al., 1986; Nishimura et al., 1995). We used brain slices submerged under artificial cerebrospinal fluid (ACSF) superfusion to avoid SD (Croning and Haddad, 1998). This setup allowed studying the detrimental effects of AGQS at their plasma concentrations separately from SD-induced injury. Using pharmacological tools combined with electrophysiology, 2PLSM imaging, and ultrastructural analyses, we investigated the involvement of VRAC, AA transport systems, and NMDA receptors in the onset of AGQS-induced neuronal injury during hypoxia in settings relevant to clinical conditions.

## Materials and Methods

### Brain slice preparation and solutions

All procedures followed National Institutes of Health guidelines for the humane care and use of laboratory animals and underwent yearly review by the Animal Care and Use Committee at the Medical College of Georgia. The mice were bred and housed in group cages in the certified animal facilities in a 12 hours light/dark cycle, with constant temperature (22 ± 1° C), and provided with food and water *ad libitum*. A total of 42 adult male and female mice at an average age of 7 months were used. The founding mice of the B6.Cg-Tg(Thy1-EGFP)MJrs/J colony were kindly provided by Dr. J. R. Sanes (Harvard University, Boston, MA) (Feng et al., 2000). The founding mice of the B6.Cg-Tg(GFAP-EGFP)1Hket colony were kindly provided by Dr. H. Kettenmann (Max Delbruck Center for Molecular Medicine, Berlin, Germany) (Nolte et al., 2001).

Brain slices (400 µm) were made according to standard protocols (Kirov et al., 2004). Mice were deeply anesthetized with isoflurane and decapitated. The brain was quickly removed and placed in cold, oxygenated (95% O_2_ / 5% CO_2_) sucrose-based artificial cerebrospinal fluid (Sucrose-ACSF) containing (in mM): 100 sucrose, 60 NaCl, 2.5 KCl, 26.2 NaHCO_3_, 1 KH_2_PO_4_, 5 MgSO_4_, 1 CaCl_2_, 11 glucose, pH 7.4, 295-297 mOsm. Transverse slices, including hippocampus, subiculum, and neocortex, were cut from the middle third of the brain using a vibrating-blade microtome (VT1000S, Leica). Immediately after sectioning, slices were recovered in a holding chamber at the interface of a standard ACSF and humidified 95% O_2_ / 5% CO_2_ atmosphere at room temperature. The standard (control) ACSF contained (in mM) 119 NaCl, 2.5 KCl, 26.2 NaHCO_3_, 1 KH_2_PO_4_, 1.3 MgSO_4_, 2.5 CaCl_2_, and 11 glucose, pH 7.4, 296–297 mOsm. D,L-2-amino-5-phosphonovaleric acid (AP5, 100 µM) was prepared as 1:500 concentrated stock solution and stored at -20° C. 4-[(2-Butyl-6,7-dichloro-2-cyclopentyl-2,3-dihydro-1-oxo-1H-inden-5-yl)oxy]butanoic acid (DCPIB, 20 µM) was prepared as 1:500 concentrated stock solution in DMSO and stored at -20° C. L-Glutamic acid γ-(p-nitroanilide) hydrochloride (GPNA, 3 mM) was prepared daily in standard ACSF at its final concentration. Stock solutions of L-alanine, glycine, and L-serine were prepared in distilled water at 1:500 final concentration and stored at -20° C. 480 µM L-alanine, 172 µM glycine, 733 µM L-glutamine, and 177 µM L-serine were added to standard ACSF daily before performing experiments. DCPIB was purchased from Tocris Bioscience. All other drugs and chemicals were obtained from Sigma-Aldrich. All drugs and AA were bath applied.

### In vitro setup for live imaging and extracellular recordings

After at least 1 h of incubation, a single slice was transferred into a submersion-type imaging/recording chamber (RC-29, 629 µL working volume, Warner Instruments) mounted on the Luigs & Neumann microscope stage. The slice was held down by an anchor (SHD-27LP/2, Warner Instruments) and superfused with oxygenated standard ACSF using a re-circulating system controlled by two peristaltic pumps (Watson-Marlow) set at the rate of 8 mL/min. The temperature was monitored by a thermistor probe and maintained at 32 ± 1°C by an in-line solution heater/cooler (CL-100, Warner Instruments) with a bipolar temperature controller (TA-29; Warner Instruments). All experiments started in control ACSF followed by a 30 min exposure to hypoxic ACSF or normoxic or hypoxic ACSF containing non-excitatory amino acids. Hypoxia was achieved by replacing O_2_ with N_2_. Reoxygenation was accomplished by returning to control ACSF bubbled with 95% O_2_ and 5% CO_2_. The field excitatory postsynaptic potentials (fEPSPs) and the slow direct current (DC) potential were recorded in the middle of the *st. radiatum* of hippocampal area CA1 with a glass microelectrode (filled with 0.9% NaCl, resistance 1–3 MΩ). fEPSPs were evoked every 15 s by stimulating Schaffer collateral pathway with current pulses (100 µs duration) delivered through a concentric bipolar microelectrode (25 μm pole separation; FHC). Signals from MultiClamp 200B amplifier (Molecular Devices) were filtered at 1 kHz and digitized at 10 kHz with Digidata 1322A interface board (Molecular Devices) to record fEPSPs. Signals were filtered at 200 Hz and digitized at 1 kHz with the MiniDigi 1 digitizer (Molecular Devices) for simultaneous continuous recording of DC potential. Electrophysiological recordings were analyzed with pClamp 10 software (Molecular Devices). Evoked synaptic responses in healthy slices had a sigmoidal input/output response function and a stable response at ½ maximal stimulation. The slope function (mV/ms) of the fEPSP was measured from the steepest 1.5 ms segment of the negative field potential to quantify synaptic strength. fEPSP slope values of every four consecutive responses were averaged to obtain a mean per minute. Data were normalized to the mean values of the responses recorded during 5 min of the baseline in standard ACSF.

### 2PLSM and image analyses

The Zeiss LSM 510 NLO META multiphoton system mounted on the motorized upright Axioscope 2FS microscope (Zeiss) was used to acquire images with an infrared-optimized 40×/0.8 NA water-immersion objective (Zeiss). The scan module was directly coupled with the Spectra-Physics Mai-Tai Ti:sapphire broadband mode-locked laser tuned to 910 nm for two-photon excitation. EGFP fluorescence was detected by internal photomultiplier tubes (PMT) of the scan module with the pinhole entirely opened. Three-dimensional (3D) time-lapse image stacks comprising ∼20 sections were taken at 1 µm increments across a 75×75 µm imaging field, using 3× optical zoom and yielding a nominal spatial resolution of 6.86 pixels/µm (12 bits per pixel, 2.51-3.2 µs pixel time). Data acquisition was controlled by LSM 510 software (Zeiss). Images of hippocampal CA1 pyramidal neurons, dendrites, and astrocytes were acquired between the recording pipette and stimulating microelectrode in the *st. radiatum* within 100 µm of the slice surface where injury from the slicing is minimal (Kirov et al., 1999; Davies et al., 2007). If shifting of the focal plane occurred due to tissue swelling, the field of focus was adjusted and re-centered before acquiring image stacks (Risher et al., 2009).

The LSM 510 Image Examiner software (Zeiss) and the Fiji image processing package distribution of ImageJ (National Instituted of Health) were used for image analysis and processing. All images were coded using the Fiji file name encrypter tool to ensure the analyses were blind to experimental conditions and experimental time points. A median filter (radius = 1) was applied to reduce photon and PMT noise. Neuronal and astroglial volume changes were analyzed using maximum intensity projections (MIPs) of image stacks because of the relatively poor axial resolution of 2PLSM (∼ 2 µm) as compared with the lateral resolution (∼ 0.4 µm). This analysis was adequate to determine relative changes in the size of neuronal and astroglial somata, which underestimated the actual volume changes assuming that cell soma volume is changing uniformly in all directions (Risher et al., 2009). Therefore, as previously described (Andrew et al., 2007; Risher et al., 2009; Risher et al., 2010), three techniques were used to detect relative changes in soma volume: (1) the cross-sectional soma area was digitally traced by hand using MIPs; (2) control and experimental MIP images were pseudocolored red and green, aligned and overlaid to reveal disparate areas that projected as red or green in contrast to yellow overlapping areas; and (3) a mask was created by tracing control MIP images and overlaid upon experimental images to reveal swollen areas from behind a mask. Small astroglial processes were not readily measurable due to the lower axial resolution. Therefore, volume changes of fine astroglial processes were not quantified. Focal swelling of dendrites which resembles beads on a string was identified by rounded regions extending beyond the initial diameter of the dendrite. Progression of dendritic swelling was assessed by quantifying the amount of dendritic beading in an imaging field following published protocols (Murphy et al., 2008; Sword et al., 2017). Briefly, the MIP of 3D image stacks was divided into 6×6 squares (12.5 x 12.5 µm), and only the squares containing dendrites were counted. The percentage of squares containing beaded dendrites was calculated.

### Intrinsic optical signal (IOS) imaging and analysis

Intrinsic optical signals were acquired as previously described (Andrew et al., 2007; Risher et al., 2011). Slices were transilluminated with a broadband halogen light source (Zeiss), and changes in light transmittance (LT) were collected with the Zeiss 2.5 x / 0.075 NA air objective with dipping cone attachment (Alexander and Nastuk, 1975). Images were captured every 5 s by the cooled AxioCam MRm CCD camera (Zeiss) controlled by AxioVision software (Zeiss). An image series revealing LT changes were created by subtracting the control image (T_cont_) from each subsequent experimental image (T_exp_) using Fiji. The difference signal (T_exp_−T_cont_) was normalized by dividing by T_cont,_ and (ΔT/T) was displayed using a pseudocolor intensity scale. Region of interest (ROI, 75 x 75 µm) corresponding to the areas of 2PLSM imaging in *st. radiatum* was created in the image series to quantify and plot the data as a percentage of the averaged digital intensity of the control ROI of that series (ΔT/T%).

### Electron microscopy and image analysis

Slices were immersed in mixed aldehydes containing 2% glutaraldehyde, 4% paraformaldehyde, 2 mM CaCl_2_, 4 mM MgSO_4_ in 0.1M cacodylate buffer at pH 7.4, rapidly fixed during 8 s of microwave irradiation, and then stored overnight in fixative at room temperature (Jensen and Harris, 1989). A small piece of tissue containing the CA1 region of the hippocampus imaged with 2PLSM was microdissected from the slice and processed for EM with osmium, uranyl acetate, dehydration, and embedding in Epon–Araldite resin using standard microwave-enhanced procedures (Kirov et al., 1999). Thin sections from the area of 2-photon imaging were cut with a diamond knife on an EM UC6 ultramicrotome (Leica), collected on pioloform-coated copper Synaptek slot grids (Electron Microscopy Sciences), and stained with uranyl acetate and lead citrate. These protocols produced well-stained and readily identifiable neuronal processes (see Fig. 1G1, G2). All chemicals for EM were from Electron Microscopy Sciences except CaCl_2_ and MgSO_4_, which were from Fisher Scientific. Ten fields were photographed randomly in each slice (total 70 fields) at a magnification of 5000x with the JEOL 1230 transmission electron microscope (JEOL) using the UltraScan 4000 camera (Gatan). The images were randomized, coded, and analyzed blind as to condition using the freely available RECONSTRUCT software (Fiala, 2005). Each image provided ∼108 µm^2^ (1080 µm^2^ per condition) to evaluate dendritic and mitochondrial swelling. Each dendritic profile (total of 1425 dendrites, 154-234 per condition) was scored as having a sign of swelling if the cytoplasm was electron lucent and microtubules were lost. Cross-sectioned dendritic mitochondria were identified, and their cross-sectional area was measured (total of 1487 mitochondria, 171-332 per condition).

**Figure 1.**
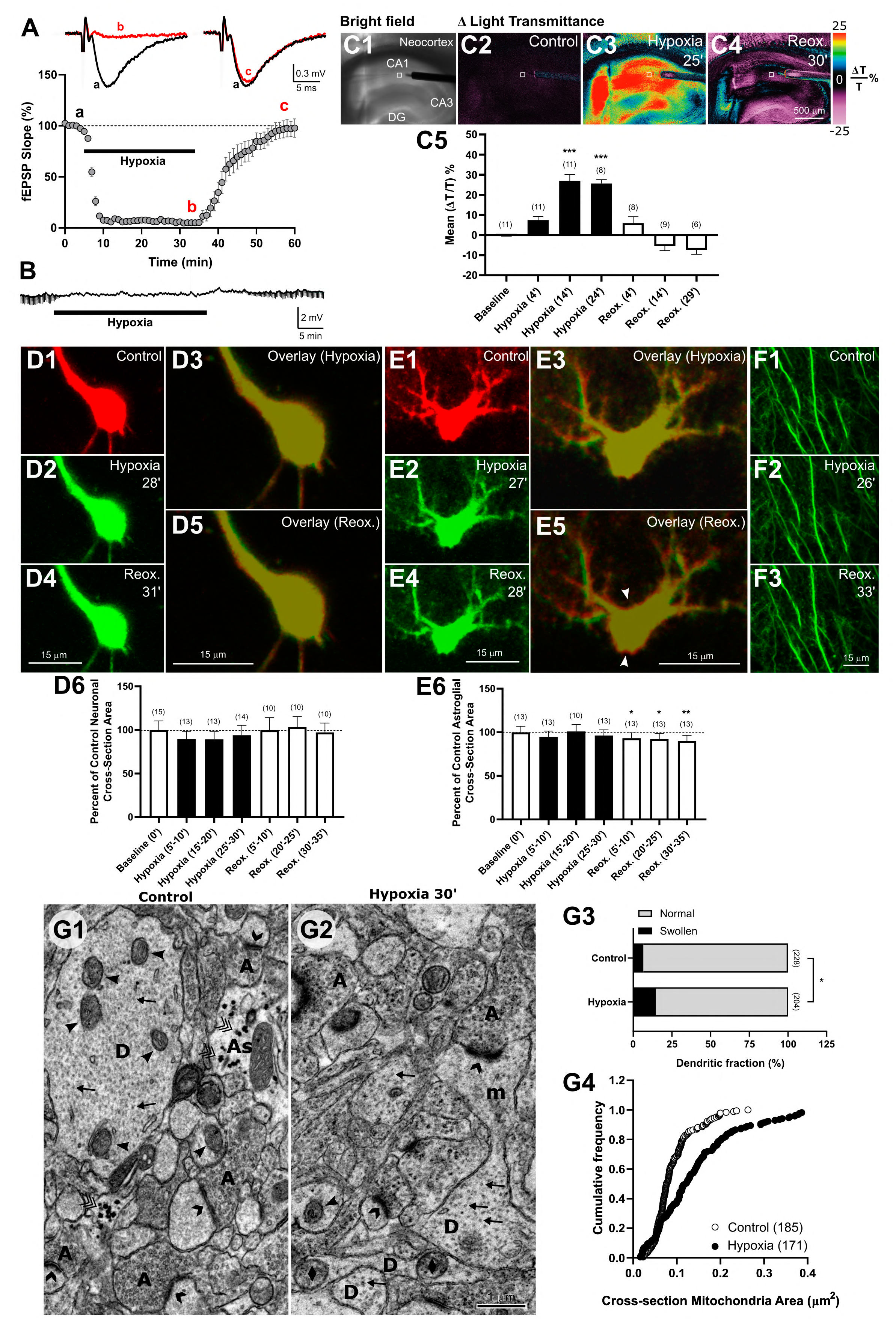
Slices from mature mice are remarkably resilient to 30 minutes of hypoxia. (**A**) Reversible loss of excitatory synaptic transmission during hypoxia; n=13 slices. Representative fEPSP traces are shown before (**a**), during hypoxia (**b**), and after reoxygenation (**c**). The stimulus artifact is truncated for clarity. (**B**) Example of continuous DC potential recording in the vicinity of 2PLSM imaging field. Small deflections at the beginning and the end of the recording represent individual fEPSPs suppressed during hypoxia. Note the lack of SD during hypoxia. (**C1-C4**) Overview of changes in LT (Δ*T/T%*) in the neocortical/hippocampal slice preparation during hypoxia and reoxygenation. Placement of stimulating electrode, glass recording pipette, and 2PLSM imaging area in *st. radiatum* (white square) are indicated in the first bright-field image (C1) acquired at the beginning of the experiment. An increase in LT during hypoxia and return to baseline levels with reoxygenation (C2-C4) are represented by pseudocolored images displayed accordingly to changes in pixel values of the bright field image (color scale: right). The white boxed zone (75 x 75 µm) corresponds to the area of 2PLSM imaging in *st. radiatum* and denotes the region used to quantify LT data shown in (C5). (**C5**) Time course of changes in LT (F_(6,30)_=29.38, P<0.0001, one-way RM ANOVA with Dunnett’s post hoc test. ***P<0.001 as compared to the baseline). Numbers in parentheses indicate the number of slices (one slice per mouse) included in the mean. (**D1**, **D2**, **D4**) MIP 2PLSM image sequence of EGFP expressing CA1 pyramidal neuron pseudocolored in red in control and green during hypoxia and reoxygenation (Reox). (**D3, D5**) Overlays displaying merged images of neuronal soma in control and 28 min of hypoxia or control and 31 min following reoxygenation reveal no changes in the neuronal volume. (**D6**) Bar graph showing no significant change in neuronal somata size during and after hypoxia (χ^2_(6)_^=7.82, P=0.25, Friedman RM ANOVA on Ranks). Numbers in parentheses indicate the number of cells included in the mean. (**E1**, **E2**, **E4**) Astrocyte morphology in control and during hypoxia and reoxygenation. (**E3, E5**) Overlays of control images and images acquired during hypoxia or reoxygenation show no change in the soma size during hypoxia but a slight decrease during reoxygenation (arrowheads). (**E6**) Time course of changes in the size of astroglial cross-section area (F_(6,54)_=3.51, P<0.01, one-way RM ANOVA with Dunnett’s post hoc test. *P<0.05, **P<0.01 as compared with the control group). Numbers in parentheses indicate the number of cells included in the mean. (**F1-F3**) 2PLSM image sequence of apical dendrites of CA1 pyramidal neurons showing no damage during and after a period of hypoxia. (**G1, G2**) Morphologically healthy neuropil from within the location of 2PLSM imaging in *st. radiatum* of the hippocampus in control and at 30 min of hypoxia. Dendrites (D) have intact cytoplasm with arrays of microtubules (arrows). Healthy synapses (chevrons) appose axonal boutons (A) filled with synaptic vesicles. An example of the well-preserved longitudinally sectioned mushroom spine (m) is also evident in the hypoxic condition. Glycogen granules (triple checkmarks) are found in large numbers in astrocytes (As) in control but not after hypoxia. Dendritic mitochondria are intact (arrowheads) in control slices but mostly swollen (diamonds) during hypoxia. (**G3**) Percentage of dendritic profiles in EM images scored as having signs of swelling was slightly increased after 30 min of hypoxia as compared to the control (χ^2_(1)_^=5.1, *P<0.05, χ^2^ test). The number of dendritic profiles analyzed in each condition is indicated to the right of each bar. (**G4**) Cumulative frequency of cross-section mitochondria area in control and hypoxic slices. Mitochondria were significantly larger in slices exposed to hypoxia (P<0.0001, K-S test).

### Statistical Analysis

Statistica (StatSoft) and SigmaPlot 14 (Systat) were used for statistical analyses. Normality was tested with the Shapiro-Wilk test. Two-tailed paired Student’s t-test and Wilcoxon’s signed-rank test, or two-tailed unpaired Student’s t-test and Mann-Whitney rank-sum test were used to compare two groups of means for parametric and nonparametric data. One-way repeated-measures analysis of variance (RM ANOVA) and Friedman RM ANOVA on Ranks followed by Dunnett’s post hoc test or two-way RM ANOVA followed by Tukey’s post hoc test were used to compare means or median values from three or more data sets. The linear regression analysis and the Pearson correlation coefficient were calculated to quantify the strength of the relationship between two variables. The slopes of the regression lines were compared by one-way analysis of covariance (ANCOVA) in case of homogeneity of regression slopes; the otherwise separate-slopes model was used. A χ^2^ test was used to analyze data arranged in contingency tables. Kolmogorov–Smirnov (K–S) test was used for statistical analysis of mitochondria size. The sample size of each experimental group is given in “Results” and “Figure legends”. The significance criterion is set at P<0.05. Data are presented as mean ± standard error of the mean (SEM).

## Results

### Submerged slices allow the analysis of brain tissue hypoxia separately from SD-triggered cellular injury

Live brain slices with preserved native tissue architecture and cellular milieu allow real-time imaging of intact neurons and astrocytes deep under the cut slice surface to observe structural changes induced by hypoxia. It is well-established that in response to transient hypoxic insult, synaptic transmission is interrupted, but neurons maintain their membrane potential, and reoxygenation restores synaptic transmission (Somjen, 2004). However, the occurrence of SD during hypoxia, as routinely observed in slices held at a gas-liquid interface, hastens rapid, irreversible neuronal injury (Müller and Somjen, 1999; Andrew et al., 2022). In contrast, SD cannot be triggered by hypoxia alone in slices submerged under normal ACSF superfusion (Croning and Haddad, 1998). Indeed, we observed, in submerged slices, a loss of hippocampal fEPSP during 30 minutes of hypoxia followed by recovery to 96.9±8.1% of baseline after 25 min of reoxygenation (t_(12)_=0.41, P=0.7, paired t-test; Fig. 1A). Importantly, no SD was recorded either electrophysiologically as an abrupt negative shift of the DC potential (n=14 slices; Fig. 1B) or optically as a sudden onset of increased LT that propagated across the slice (n=11 slices; Fig. 1C) (Andrew et al., 2007; Risher et al., 2011). Indeed, LT slowly and simultaneously elevated throughout the hippocampus peaking at ∼27% at 14 min after starting hypoxia (P<0.0001, one-way RM ANOVA) and returning to baseline during reoxygenation.

Since an increase in LT has been shown to be reflecting the swelling of brain cells and has also been hypothesized to be due to the swelling of intracellular organelles (e.g., mitochondria) (Jarvis et al., 1999), we used 2PLSM to monitor volume changes of EGFP-expressing neurons and astrocytes and EM to measure mitochondrial swelling. Hippocampal pyramidal CA1 neurons steadfastly maintained their volume during hypoxia and reoxygenation (Fig. 1D1-D6; P=0.25, Friedman RM ANOVA on Ranks). Astroglial soma size was also not affected by hypoxia but slightly decreased by 10.1±6.6% at the end of 30 min of reoxygenation (Fig. 1E1-E6; P=0.005, one-way RM ANOVA). The dendritic structure was preserved during hypoxia (Fig. 1F1-F3), ensuing recovery of synaptic transmission during reoxygenation. The amount of dendritic beading in the imaging field is considered a reliable indicator of neuronal injury and cytotoxic edema (Hsu and Buzsáki, 1993; Risher et al., 2010; Kirov et al., 2020). Indeed, there was no change in the rate of dendritic beading, which was never over 5% at any time point, indicating that hypoxia was not causing damage to the synaptic circuitry (n=7 imaging fields, χ^2_(6)_^ =6.0, P=0.42, Friedman RM ANOVA on Ranks). This finding at the level of the light microscopy was corroborated with ultrastructural analyses that reveal largely intact neuropil with healthy synapses after 30 min hypoxia as compared to the control tissue (Fig. 1G1, G2). However, there was a slight increase in dendritic profiles with signs of swelling from ∼7% in control to ∼15% at 30 min of hypoxia (Fig. 1G3, P=0.02, χ^2^ test). The average cross-section mitochondria area also increased by 57.2±8.3% during hypoxia (P<0.001, Mann-Whitney rank sum test), revealing mitochondrial swelling (Fig. 1G4).

Together, these analyses conducted in the CA1 area of the hippocampus indicate that after 30 minutes of hypoxia and despite significant swelling of dendritic mitochondria, slices remain relatively healthy with unimpaired synaptic responses or without undergoing SD or experiencing cytotoxic astroglial and neuronal edema and dendritic injury.

### AGQS induce profound neuronal damage and persistent astroglial swelling during hypoxia

To determine if AGQS cause any harm when combined with hypoxia, we applied these AA at their plasma concentration to hypoxic ACSF. fEPSPs suffered irreversible silencing that remained during the washout and reoxygenation period, with fEPSPs returning only to 14.6±8.9% of the baseline (Fig. 2A, P=0.001, Wilcoxon signed-rank test). SD was never detected in any of our electrophysiological recordings (n=12 slices, Fig. 2B), indicating that the permanent loss of synaptic transmission cannot be attributed to the occurrence of SD. IOS imaging also confirmed that AGQS during hypoxia were not causing SD, which would be detectable as a spreading wave of increased LT. The LT raised gradually throughout the hippocampus to ∼40% at 14 min of the hypoxia (Fig. 2C1-C5, P<0.001, one-way RM ANOVA). At this time point, the increment in LT was significantly larger than during hypoxia alone (t_(18)_=2.87, P=0.01, t-test), suggesting evolving cellular swelling in the presence of AGQS. The LT started to decrease before the end of hypoxia and washout of AGQS (Fig. 2C5). This decrease in LT likely indicated the onset of dendritic beading, which scatters light, lowering LT in swollen tissue (Jarvis et al., 1999; Andrew et al., 2007; Douglas et al., 2011). Next, we used 2PLSM to monitor in real-time structural changes implied by changes in LT.

**Figure 2.**
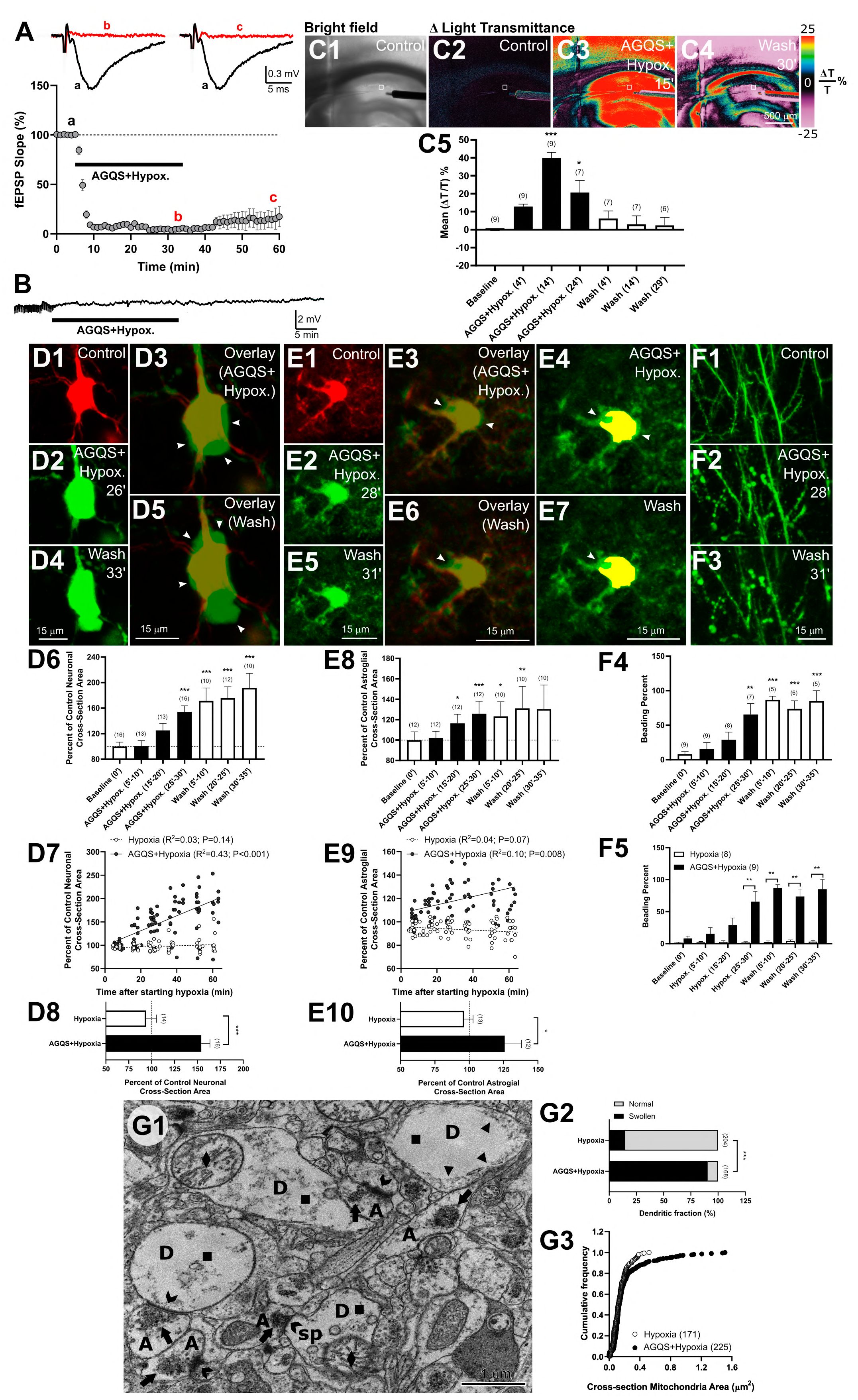
Irreversible damage to neurons and enduring astroglial swelling induced by AGQS during hypoxia. (**A**) Permanent loss of fEPSPs after hypoxia in the presence of AGQS; n=12 slices. Examples of fEPSP traces recorded in control (**a**), at the end of AGQS application during hypoxia (**b**), and washout during reoxygenation (**c**). (**B**) DC recording from a representative experiment shows that SD was absent during hypoxia and recovery. Small deflections at the beginning of the recording denote fEPSPs that permanently disappear during and after hypoxia in the presence of AGQS. (**C1**) Representative bright-field photograph of a brain slice illustrating the position of glass recording pipette, stimulating electrode, and area of 2PLSM imaging in *st. radiatum* (white square). (**C2-C4**) Pseudocolored images of changes in LT during the time course of the experiment. The white square denotes the area of 2PLSM imaging. (**C5**) Changes in LT quantified over the area of 2PLSM (F_(6,18)_=10.76, P< 0.0001, one-way RM ANOVA with Dunnett’s post hoc test. *P<0.05, ***P<0.001 as compared with the baseline). (**D1-D5**) CA1 neurons swell in the presence of AGQS during hypoxia. Pseudocolored in red control image of neuronal soma is overlaid with pseudocolored in green neuronal soma during 26 min of exposure to hypoxic ACSF containing AGQS (D3) and during 33 min of washout and reoxygenation (D5). Arrowheads are pointing to swollen areas. (**D6**) Bar graph illustrating the significant increase in neuronal cross-section somata area during hypoxia in the presence of AGQS and after washout and reoxygenation (F_(6,54)_=21.27, P< 0.0001, one-way RM ANOVA with Dunnett’s post hoc test. ***P<0.001 represent a significant difference from control). (**D7**) Regression line shows a strong positive correlation between neuronal somata size and the duration of the hypoxia in the presence of AGQS (n=16 neurons). Without AGQS, there was no significant relationship between somata size and hypoxia duration (n=15 cells). Neurons were significantly larger when exposed to AGQS during hypoxia (F_(1,154)_=126.55, P<0.0001, one-way ANCOVA controlling for the effect of time after the start of hypoxia). (**D8**) Changes in neuronal cross-section somata area in the last 5 minutes of hypoxia alone or with AGQS (***P<0.001, Mann-Whitney Rank Sum test). (**E1, E2, E5**) MIP image sequence of astrocyte pseudocolored red in control and green for two experimental conditions. (**E3**, **E6**) Overlaid images illustrating astroglial swelling with arrowheads pointing to swollen areas. (**E4**, **E7**) Yellow mask representing the control soma area is placed over the same soma following subsequent treatments to reveal swelling (arrowheads). (**E8**) Bar graph showing a significant increase in astroglial cross-section somata area during and after AGQS with hypoxia. (χ^2_(6)_^=31.57, P<0.001, Friedman RM ANOVA on Ranks with Dunnett’s post hoc test. *P<0.05, **P<0.01, ***P<0.001 represent a significant difference from control). (**E9**) Regression lines showed a positive correlation between astroglial swelling and time after hypoxia with AGQS (n=12 astrocytes), while there was no such significant correlation in the astroglial somata size during hypoxia without AGQS (n=13 cells). Astrocytes were significantly larger when exposed to AGQS during hypoxia (F_(1, 143)_=6.11, P<0.05, separate-slopes model controlling for the effect of time after the start of hypoxia). (**E10**) Changes in astroglial cross-section somata area in the last 5 minutes of hypoxia with and without AGQS (*P<0.05; unpaired t-test). (**F1**-**F3**) Sequence of MIP images revealing persistent dendritic beading upon exposure to AGQS during hypoxia and washout. (**F4**) Quantification of dendritic beading showing a significant increase during hypoxia with AGQS and during washout (F_(6,24)_=10.27, P<0.0001, one way RM ANOVA followed by Dunnett’s post hoc test. **P<0.01, ***P<0.001 as compared to the control). The number of analyzed mice is indicated above each bar. (**F5**) Hypoxia alone (n=8 mice) does not induce dendritic beading as compared with hypoxia in the presence of AGQS (n=9 mice) (F_(1,7)_=23.88, P<0.001, two way RM ANOVA with Tukey’s HSD post hoc test. **P<0.01 represent significant differences between groups at each time point). (**G1**) Ultrastructural components of damaged synaptic circuitry 30 min after exposure to hypoxic ACSF containing AGQS. Severely swollen dendrites (D) were without microtubules and had a watery cytoplasm (squares), thin cytoplasmic rims (triangles), and swollen mitochondria (diamonds). Some dendrites had collapsed spines (Sp), or spines were overwhelmed by swelling, but synapses (chevrons) were present in the neuropil. Axonal boutons (A) were severely swollen with clumped synaptic vesicles either next to release sites or at the center of profiles (arrows). (**G2**) The fraction of swollen dendritic profiles in electron micrographs was significantly higher when AGQS were present during hypoxia (χ^2_(1)_^ =111.1, ***P<0.001, χ^2^ test). (**G3**) Mitochondria were significantly larger during exposure to hypoxia in the presence of AGQS than in hypoxia alone (P<0.02, K-S test).

2PLSM showed potent and irreversible neuronal swelling in the presence of AGQS during hypoxia which peaked at 191.6±22.9% at the end of the washout period (Fig. 2D1-D6; P<0.0001, one-way RM ANOVA). The linear regression analysis revealed a strong relationship between neuronal somata size and time passed after the beginning of hypoxia with AGQS (Fig. 2D7; P<0.001, Pearson correlation). In contrast, there was no such relationship when neurons were exposed to hypoxia without amino acids (Fig. 2D7; P=0.14, Pearson correlation). One-way ANCOVA revealed that neurons were significantly enlarged in the presence of AGQS (P<0.0001). Indeed, the difference in size was clearly noticeable in the last 5 minutes of hypoxia with AGQS than without (Fig. 2D8; P<0.001, Mann-Whitney rank sum test).

Similarly, 2PLSM revealed significant and irreversible astroglial swelling when AGQS were applied during hypoxia (Fig. 2E1-E7). Quantification has shown that the astroglial cross-section somata area remained significantly enlarged by ∼31% relative to the baseline during the 30-minute washout period (P<0.001, Friedman RM ANOVA on Ranks). In contrast to hypoxia alone, when there was no significant positive correlation between astroglial somata size and time after the beginning of hypoxia (Fig. 2E9; P=0.07, Pearson correlation), this correlation was significant in the presence of AGQS (P=0.008, Pearson correlation). Accordingly, astrocytes were significantly bigger in slices exposed to AGQS during hypoxia than those exposed only to hypoxia without AA (Fig. 2E9; P=0.015, separate-slopes model). The difference in the astroglial soma size was apparent in the last 5 minutes of exposure to hypoxia with AGQS than without (Fig. 2E10; t_(23)_=2.19, P=0.04, t-test).

2PLSM revealed remarkable dendritic beading during hypoxia with AGQS (Fig. 2F1-F3). Quantification showed that beading was permanent (Fig. 2F4; P<0.0001, one-way RM ANOVA), pointing to the terminal damage to the synaptic circuitry, which was accompanied by the irreversible loss of synaptic transmission (Fig. 2A). Once again, there was no dendritic beading caused by hypoxia alone (Fig. 1F1-F3 and Fig. 2F5). Significant dendritic beading was detected in the last 5 min of hypoxia with AGQS (Fig. 2F4, F5), likely causing the decrease in the LT at the end of the hypoxic period (Fig. 2C5).

Structural changes that were first observed on living dendrites with 2PLSM were supported by EM analyses (Fig. 2G1-G3). Dendrites showed signs of severe beading as evidenced by watery cytoplasm, thin cytoplasmic rims, and the disappearance of microtubules (Fig. 2G1) (Kirov et al., 2020). About 90% of dendritic profiles showed signs of swelling after 30 min of hypoxia with AGQS, a significant increase from about only 15% during hypoxia alone (Fig. 2G2, P<0.001, χ^2^ test). Likewise, axonal boutons were grossly swollen with clumped synaptic vesicles, but large numbers of them remained in the boutons. Clumped vesicles indicate dysfunctional synaptic contacts, nonetheless some vesicular release may occur at the onset of swelling (Kovalenko et al., 2006). The average cross-section mitochondria area was 140.4±18.6% bigger during hypoxia with AGQS than in control (P<0.001, Mann-Whitney rank sum test). This increase in mitochondria size corresponded to an additional swelling by 52.9±11.8% when compared to hypoxia without AGQS (P=0.13, Mann-Whitney rank sum test) (Fig. 2G3). Notably, the largest mitochondria cross-sectional area during hypoxia with AGQS was 1.51 µm^2^, which was 573.3% larger than the largest mitochondria cross-sectional area in control (0.26 µm^2^) and 290.6% larger than the maximum size detected during hypoxia (0.52 µm^2^). It is conceivable that such dramatic swelling in the presence of AA during hypoxia resulted in the burst of some of these organelles. Accordingly, 51.8% of cross-sectioned dendritic profiles were without mitochondria in the presence of AGQS and hypoxia as compared to only 15.3% in the control condition (χ^2_(1)_^=30.79, P<0.001, χ^2^ test) and 27.7% in hypoxia (χ^2_(1)_^=9.42, P=0.002, χ^2^ test).

Together, these results indicate extensive irreversible tissue damage caused by a mixture of AGQS during hypoxia without the occurrence of SD. Under hypoxic conditions, AGQS induce astroglial swelling and a significant increase in neuronal soma size with severe dendritic beading, which translates into a terminal silencing of synaptic transmission.

### AGQS cause swelling of astrocytes, but not hippocampal pyramidal neurons

To examine if the presence of AGQS exerts any adverse physiological effects under normoxic conditions, we assessed synaptic transmission and changes in LT before, during, and after the application of AGQS (Fig. 3A, B). Electrophysiological recordings revealed significant potentiation of fEPSPs by 13.3±3.25% at the end of the AGQS application period (Fig. 3A; P=0.001, Wilcoxon signed-rank test). The LT transiently raised, indicating tissue swelling, and peaked at an 11.2±3.8% increase at the beginning of the washout period (Fig. 3B; P<0.001, Friedman RM ANOVA on Ranks). The swelling was already detectable at 14 min of normoxic AGQS application (6.6±0.9% increase in the LT; P<0.05, Dunnett’s post hoc test). However, the LT increment was significantly milder than during the same time point with AGQS in hypoxia (39.9±9.4% increase in the LT; Fig. 2C5) (P<0.001, Mann-Whitney rank sum test when comparing the LT raise in slices exposed to normoxic vs. hypoxic AGQS mixtures).

**Figure 3.**
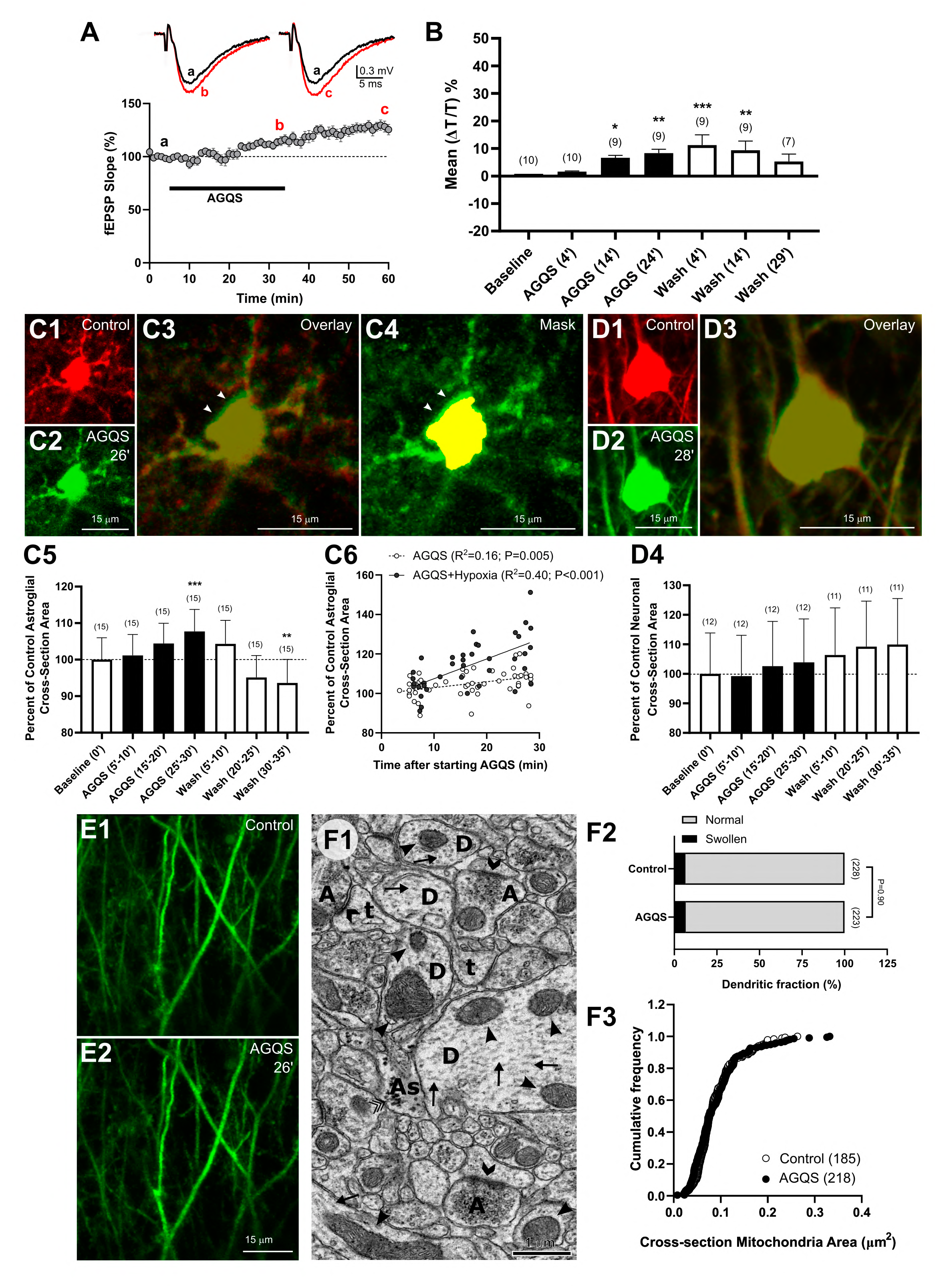
Astrocytes but not neurons swell in the presence of AGQS. (**A**) AGQS induces a significant potentiation of evoked field potentials; n=13 slices. Representative fEPSP recordings in control (**a**), at the end of AGQS application (**b**), and during washout (**c**). (**B**) Transient increase in LT during AGQS application provided evidence of slice swelling and recovery when AGQS were washed out. (χ^2_(6)_^ = 24.86, P<0.001, Friedman RM ANOVA on Ranks with Dunnett’s post hoc test. *P<0.05, **P<0.01, ***P<0.001 as compared to the baseline). (**C1**-**C3**) Pseudocolored MIP images of astrocyte in control (red) and during AGQS application (green) are overlayed to illustrate an increase in astroglial volume (arrowheads). (**C4**) The yellow mask created from the control image of the astrocyte is used to reveal swollen areas (arrowheads) of the astrocyte exposed to AGQS. (**C5**) Quantification of significant astroglial swelling induced by AGQS (F_(6,90)_=13.42, P<0.0001, one-way RM ANOVA with Dunnett’s post hoc test. **P<0.01, ***P<0.001 as compared to the control). (**C6**) Regression line shows a positive correlation between astroglial somata size and time of exposure to AGQS during normoxia (n=15 astrocytes) or hypoxia (n=12). Astrocytes were significantly larger when exposed to AGQS during hypoxia as compared to AGQS in normoxic conditions (F_(1,84)_=26.10, P<0.0001, one-way ANCOVA controlling for the effect of time after start of exposure to AGQS). (**D1**-**D3**) Overlay showing merged images of CA1 pyramidal neuron captured in control (red) and during exposure to AGQS (green) reveals no increase in soma volume at 28 min of AGQS application. (**D4**) Quantification of the size of neuronal somata in control and AGQS during normoxia confirms no significant changes (F_(6,60)_=1.57, P=0.17, one-way RM ANOVA controlling for the duration of AGQS exposure). (**E1**-**E2**) Representative MIP images of apical dendrites of CA1 pyramidal neurons displaying no effect of AGQS on dendritic morphology. (**F1**) There were no changes in the ultrastructure of the neuropil of the normoxic slice exposed to AGQS for 30 min. Morphologically healthy dendrites (D) contained intact cytoplasm with arrays of microtubules (arrows) and undamaged mitochondrial organelles (arrowheads). Well-preserved longitudinally sectioned thin (t) dendritic spines are also evident. Synapses (chevrons) and axonal boutons (A) filled with synaptic vesicles are well preserved. Astrocytes (As) contained numerous glycogen granules (triple checkmarks). (**F2**) Percentage of dendritic profiles in EM images with signs of swelling was similar between control and AGQS conditions (χ^2_(1)_^=0.02, P=0.9, χ^2^ test). (**F3**) The size of dendritic mitochondria was similar between control and AGQS conditions (P=0.12, K-S test).

The 2PLSM revealed that the transient increase in the LT caused by AGQS during normoxia was likely due to the transient swelling of astrocytes (Fig. 3 C1-C5; P<0.0001, one-way RM ANOVA) as neuronal volume was not affected (Fig. 3D1-D4; P=0.17, one-way RM ANOVA). Indeed, the linear regression analysis has shown a positive correlation between astroglial somata size, and the time slices were exposed to AGQS during normoxia (Fig. 3C6; P=0.005, Pearson correlation). Nevertheless, the comparison of regression lines indicated that astrocytes were further swollen when exposed to AGQS during hypoxia (Fig. 3C6; P<0.0001, one-way ANCOVA).

Additionally, none of the six dendritic fields analyzed in the different slices from six animals showed any sign of beading from exposure to the AGQS mixture (Fig. 3E1, E2). Accordingly, EM analyses revealed well-preserved neuropil with intact dendrites, axons, and synapses (Fig. 3F1). Indeed, AGQS did not cause any increase in the proportion of the dendritic profiles with signs of swelling compared to the control (Fig. 3F2; P=0.90, χ^2^ test). Dendritic mitochondria have also remained intact and similar in their size to the control mitochondria (Fig. 3F3; P=0.12, K-S test).

Our results have shown that astrocytes swell in the presence of AGQS, but neurons and their dendrites remain intact. Conceivably, the lack of neuronal soma and dendritic mitochondria swelling could explain a smaller increase in the LT in slices during AGQS application in normoxic than in hypoxic conditions.

### Hippocampal neurons and astrocytes are resistant to AGS or glutamine during hypoxia

Since L-alanine, glycine, and L-serine in the AGQS mixture are NMDAR co-agonists (Pace et al., 1992), and glutamine can be metabolically converted into glutamate in neurons an astrocytes (Bak et al., 2006), we tested if the mixture of AGS or glutamine alone can evoke similar damage to the synaptic circuitry during hypoxia as four AA combined.

The fEPSP loss induced by hypoxia with AGS was entirely reversible, recovering to 115.0±8.7% of baseline after 25 minutes of washout and reoxygenation (Fig. 4A; P=0.11, Wilcoxon signed-rank test). The exposure of the slices to glutamine during hypoxia also did not significantly affect the recovery of synaptic transmission during washout and reoxygenation (Fig. 4B; P=0.17, paired t-test). No SD was detected in any 14 slices superfused with AGS or glutamine during hypoxia.

**Figure 4.**
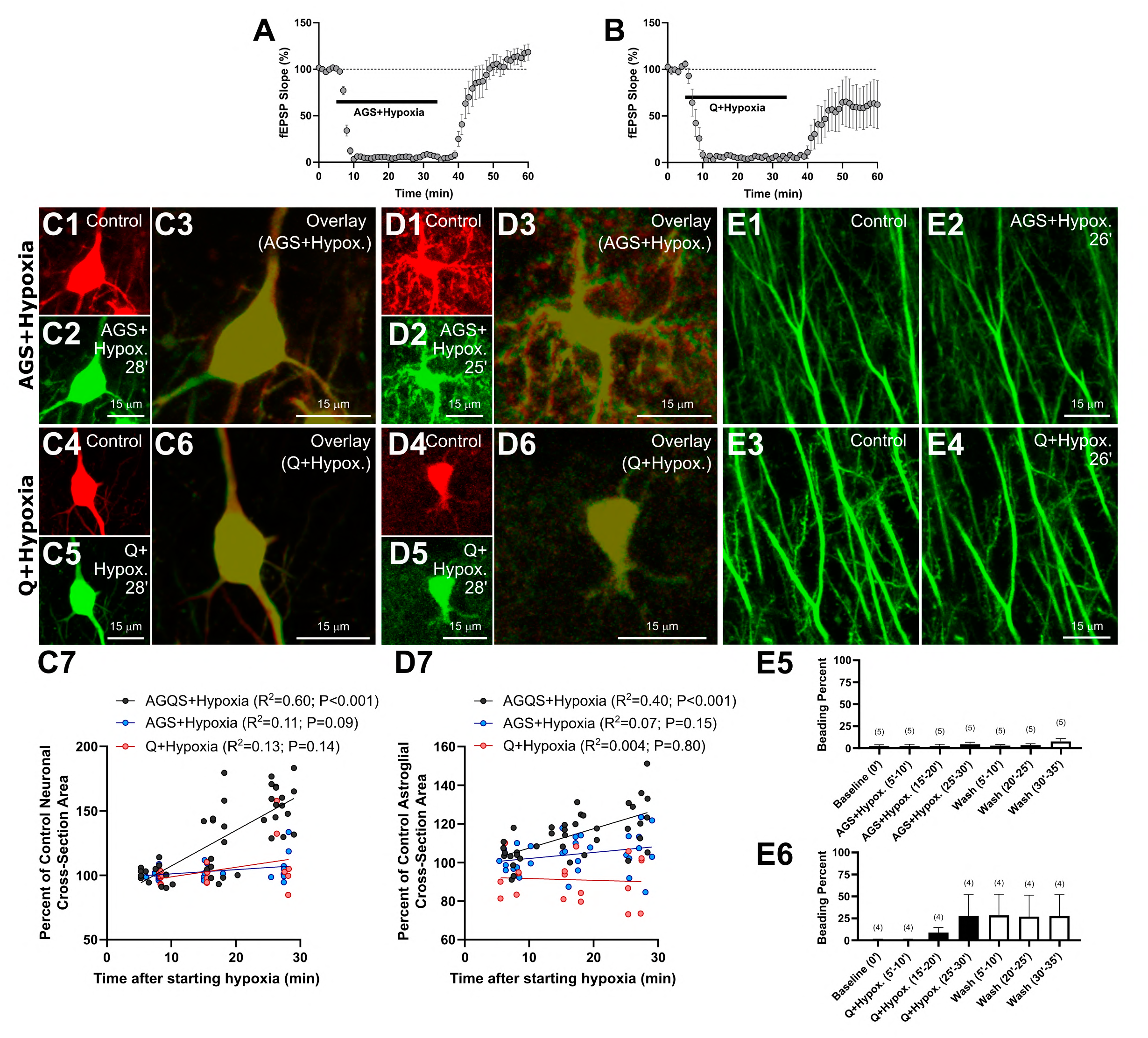
Neurons and astrocytes withstand exposure to AGS or glutamine (Q) during 30 min of the hypoxic period. (**A**) Reversible loss of fEPSPs during hypoxia in the presence of AGS; n=8 slices. (**B**) Excitatory synaptic transmission was lost during hypoxia in the presence of glutamine (Q) but recovered during washout and reoxygenation to 61.6±24.1% of the baseline (n=6 slices; t_(5)_=1.61, P=0.17, paired t-test). (**C1**-**C2, C4-C5**) Color-coded images of neuronal soma in control (red) and during hypoxia (green) in the presence of either AGS or glutamine (Q). (**C3**, **C6**) Corresponding overlays show no changes in neuronal soma volume. (**C7**) There was no correlation between neuronal somata size and time after the start of hypoxia with either AGS (n=9 cells) or glutamine (Q; n=6) as compared to AGQS (n=16). Neurons were swollen when exposed to AGQS during hypoxia in comparison to AGS or glutamine (F_(2,85)_=18.70, P<0.0001, one-way ANCOVA controlling for the effect of time after start of hypoxia). The astroglial soma area imaged in control (**D1** and **D4**) overlaps entirely with the astroglial soma area imaged in hypoxia with AGS (**D2**) or glutamine (Q; **D5**), illustrating no changes in soma size as seen in corresponding color-coded overlays (**D3** and **D6**). (**D7**) In contrast to AGQS (n=12 astrocytes), regression lines show no correlation between astroglial somata size and duration of hypoxia with AGS (n=9 cells) or glutamine (Q; n=5). Astrocytes swell when exposed to AGQS during hypoxia in comparison to AGS or glutamine (F_(2,82)_=29.03, P<0.0001, one-way ANCOVA controlling for the effect of hypoxia duration). (**E1-E2** and **E3-E4**) Paired MIP images of dendrites of CA1 pyramidal neurons illustrate no damage to the dendrites during hypoxia in the presence of AGS or glutamine (Q) as confirmed by quantitative analyses for AGS (**E5**) (χ^2_(6)_^=9.46, P=0.15, Friedman RM ANOVA on Ranks) or glutamine (**E6**) (χ^2_(6)_^=10.38, P=0.11, Friedman RM ANOVA on Ranks).

The neuronal soma volume was unaffected by the presence of AGS or glutamine during hypoxia (Fig. 4C1-C6). The linear regression analyses in Fig. 4C7 revealed no significant relationship between neuronal somata size and time after hypoxia with AGS (P=0.09, Pearson correlation) or glutamine (P=0.14, Pearson correlation). The slopes of regression lines for AGS and glutamine differ significantly from the slope of the regression line for AGQS (Fig. 4C7; P<0.0001, one-way ANCOVA), indicating a continuous increase in somata size after exposure to AGQS during hypoxia as compared to AGS or glutamine.

Astrocytes in slices exposed to AGS or glutamine were not swollen during hypoxia (Fig. 4D1-D6). Astroglial cells maintained their volume (Fig. 4D7) throughout hypoxia with AGS (P=0.15, Pearson correlation) or glutamine (P=0.80, Pearson correlation). This lack of changes in astroglial volume was in contrast to continuing astroglial volume increase during hypoxia with AGQS (Fig. 4D7; P<0.0001, one-way ANCOVA).

Dendrites were intact after 30 minutes of exposure to AGS or glutamine during hypoxia (Fig. 4E1-E4). Quantification of five dendritic fields from five slices from different mice has shown no signs of dendritic beading during hypoxia with AGS and washout with reoxygenation (Fig. 4E5; P=0.15, Friedman RM ANOVA on Ranks). Likewise, there was no significant dendritic damage detected in four slices exposed to hypoxia in the presence of glutamine (Fig. 4E6; P = 0.11, Friedman RM ANOVA on Ranks).

These results indicate that the combination of all four AGQS AA is necessary to induce profound and irreversible damage to nervous tissue during hypoxia. Exposure to the mixture of AGS or glutamine alone was innocuous and did not cause any harm.

### Blockade of NMDAR prevents hypoxic damage induced by AGQS

Recently we have shown that the irreversible loss of synaptic potentials caused by a mixture of seven non-excitatory amino acids (AGQHSTU) during hypoxia is NMDAR-dependent (Álvarez-Merz et al., 2021). Here, we investigated if AP5, a selective and competitive antagonist of NMDAR, could prevent AGQS-induced neuronal damage during hypoxia.

DL-AP5 (100 µM) precluded irreversible loss of fEPSPs resulting from 30 min hypoxia with AGQS (Fig. 5A). fEPSPs recovered, reaching a slight potentiation of 36.3±14.5% at the end of the washout and reoxygenation period (P=0.04, Wilcoxon signed-rank test). Accordingly, neuronal swelling during hypoxia with AGQS was eliminated when NMDAR were blocked (Fig. 5B1-B4; P=0.6, one-way RM ANOVA) and neuronal somata were significantly smaller in the presence of AP5 (Fig. 5B5; P=0.02, separate-slopes model). Moreover, there was no significant relationship between neuronal somata size and time during hypoxia with AGQS when AP5 was present (Fig. 5B5; P=0.73, Pearson correlation).

**Figure 5.**
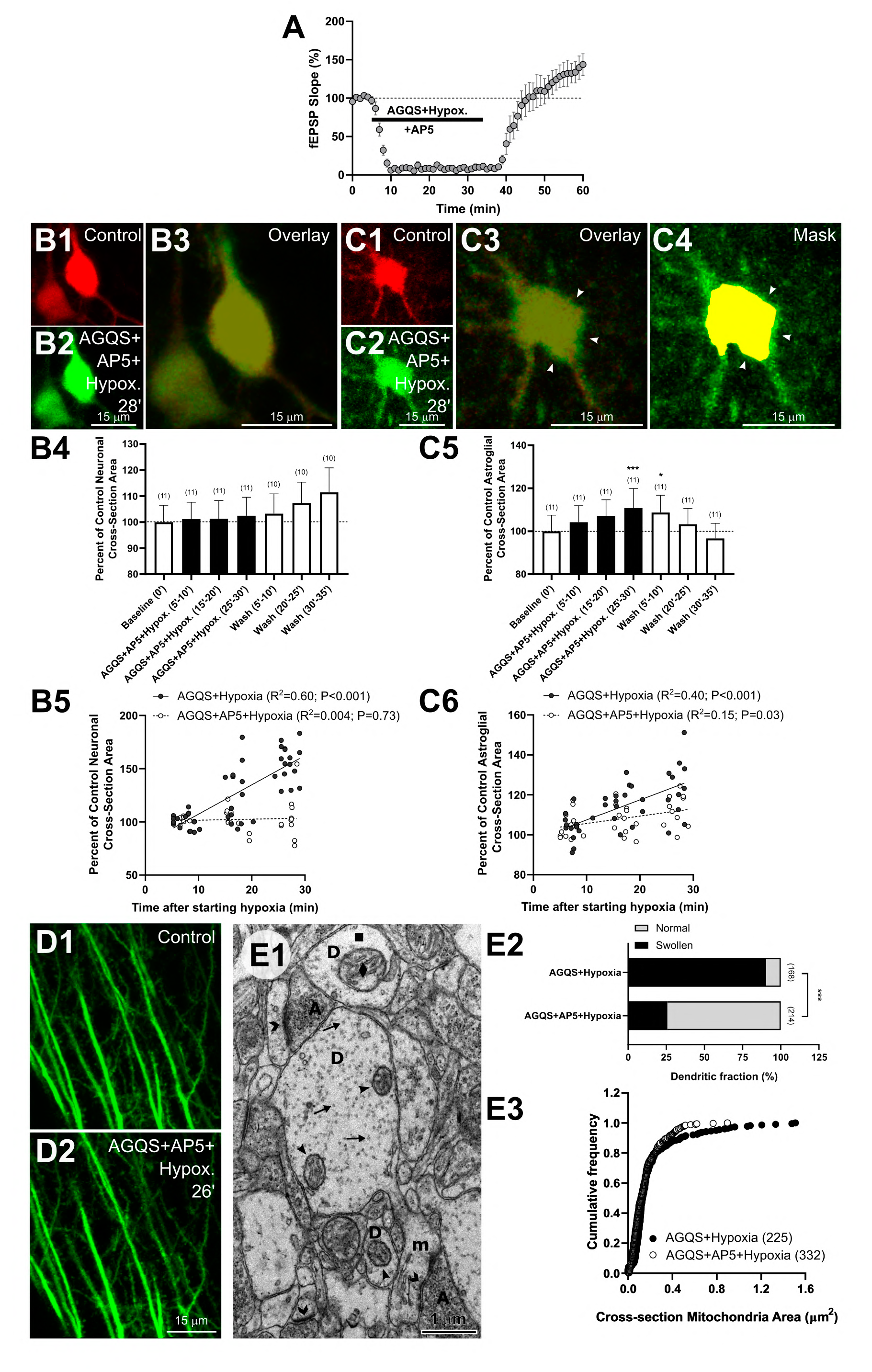
AGQS-induced hypoxic neuronal damage is NMDAR-dependent. (**A**) Synaptic transmission, transiently abolished during hypoxia in the presence of AGQS and AP5, was potentiated after washout and reoxygenation (P<0.05, Wilcoxon signed-rank test, n=10 slices). (**B1**-**B3**) Blocking NMDAR with AP5 prevented neuronal swelling elicited by hypoxia in the presence of AGQS, as revealed by overlaid images of neuronal soma in control and hypoxic conditions. (**B4**) Quantification confirmed no changes in the size of neuronal cross-section area during hypoxia in the presence of AGQS and AP5 (F_(6,54)_=0.77, P=0.6, one-way RM ANOVA). (**B5**) Similarly, there was no correlation between neuronal somata size and duration of hypoxia when AP5 was present together with AGQS (n=11 neurons). Neurons were not swollen when NMDAR were blocked during hypoxia with AGQS (F_(1,73)_=5.45, P<0.05, separate-slopes model controlling for the effect of time after the start of hypoxia). (**C1**-**C3**) Color-coded overlay of astroglial soma in control and during hypoxia with AGQS and AP5 shows swelling (arrowheads). (**C4**) The yellow mask corresponding to the astroglial soma area in control is placed over the same soma during hypoxia with AGQS and AP5. Arrows point to swollen areas. (**C5**) Time-course of astroglial swelling during treatment with AGQS and AP5 under hypoxic conditions and washout and reoxygenation (F_(6,60)_=6.73, P<0.0001, one-way RM ANOVA with Dunnett’s post hoc test. *P<0.05, ***P<0.001 relative to the control). (**C6**) Regression line shows a positive correlation between astroglial somata size and duration of hypoxia with AP5 and AGQS (n=11 cells). Astrocytes were smaller during hypoxia with AP5 and AGQS than with hypoxia and AGQS (F_(1,67)_=7.11, P<0.01, one-way ANCOVA controlling for the effect of time after the start of hypoxia). (**D1**-**D2**) Paired images of dendrites in control and during subsequent treatment with AGQS and AP5 during hypoxia reveal thorough preservation of the dendritic structure. (**E1**) Neuropil of area CA1 after 30 min of hypoxia in the presence of AGQS and AP5. The majority of dendrites (D) contained intact cytoplasm with microtubules (arrows) and intact mitochondria (arrowheads). However, some mitochondrial organelles were swollen (diamonds). An example of the well-preserved longitudinally sectioned mushroom spine (m) with postsynaptic density (chevron) apposed by axonal bouton (A) full of synaptic vesicles is clearly visible. Some dendrites had signs of swelling with a watery cytoplasm (squares) and distorted microtubule arrays. (**E2**) Significantly smaller fraction of dendrites in electron micrographs had signs of swelling when NMDAR were blocked with AP5 during hypoxic periods with AGQS than without AP5 (χ^2_(1)_^ = 4.65, ***P<0.001, χ^2^ test). (**E3**) Dendritic mitochondria were significantly smaller when NMDAR were blocked during hypoxia with AGQS (P<0.001, K-S test).

As expected, astrocytes were swollen during hypoxia with AGQS and the NMDAR antagonist, and the swelling was reversible upon the washout and reoxygenation (Fig. 5C1-C5; P<0.0001, one-way RM ANOVA). The astroglial volume continued to increase during hypoxia with AGQS and AP5 (Fig. 5C6; P=0.03, Pearson correlation). However, the rate of the increase was slower than with AGQS alone (Fig. 5C6; P=0.01, one-way ANCOVA), probably due to the blockade of functional NMDAR expressed in astrocytes (Schipke et al., 2001; Verkhratsky and Chvátal, 2020).

Importantly, blockade of NMDAR with AP5 stopped dendritic beading caused by AGQS during hypoxia (Fig. 5D1-D2). Quantification of seven dendritic fields showed that the dendrites were not beaded throughout the experiment (F_(6,24)_=1.0, P=0.45, one-way RM ANOVA; data not shown). Similarly, EM revealed well-preserved dendrites with healthy synapses (Fig. 5E1). The majority of dendrites contained intact cytoplasm as the fraction of dendrites with signs of swelling decreased from ∼90% during hypoxia with AGQS to ∼26% when AP5 was added (Fig. 5E2; P<0.001, χ^2^ test). Dendritic mitochondria were less swollen with AP5 during hypoxia and AGQS than without AP5 (Fig. 5E3, P<0.0001, K-S test). Indeed, mitochondria were swollen to the same degree as during hypoxia alone (P=0.95, Mann-Whitney rank sum test; data not shown).

Therefore, the results reveal that activation of NMDAR is implicated in the irreversible neuronal damage induced by AGQS during hypoxia.

### VRAC are involved in AGQS-induced cellular injury during hypoxia

NMDAR activation is implicated in excitotoxicity and neuronal damage (Choi et al., 1988; Vander Jagt et al., 2008; Yan et al., 2020). We hypothesized that one of the possible sources of excitotoxins in our experiments is their release through VRAC (Rosenberg et al., 2010; Hyzinski-García et al., 2014; Mongin, 2016) that may be opened during astroglial swelling induced by AGQS. Accordingly, blockade of VRAC with DCPIB (20 µM) abolished neuronal swelling caused by AGQS during hypoxia (Fig. 6A1-A4; P=0.99, one-way RM ANOVA). Regression analyses revealed that neurons steadfastly maintained their volume during hypoxia with AGQS and DCPIB (Fig. 6A5; n=10 neurons, P=0.96, Pearson correlation). Surprisingly, astroglial swelling was not readily detectable during hypoxia with AGQS and blocked VRAC (Fig. 6B1-B4; P=0.03, one-way RM ANOVA, but Dunnett’s post hoc tests used to compare to the control values were not significant). On the other hand, there was a positive correlation between astroglial soma size and time after hypoxia (Fig. 6B5; P=0.004, Pearson correlation), indicating some astrocytic swelling in the presence of DCPIB. Yet, this swelling was significantly less pronounced than without DCPIB (Fig. 6B5; P<0.0001, one-way ANCOVA).

**Figure 6.**
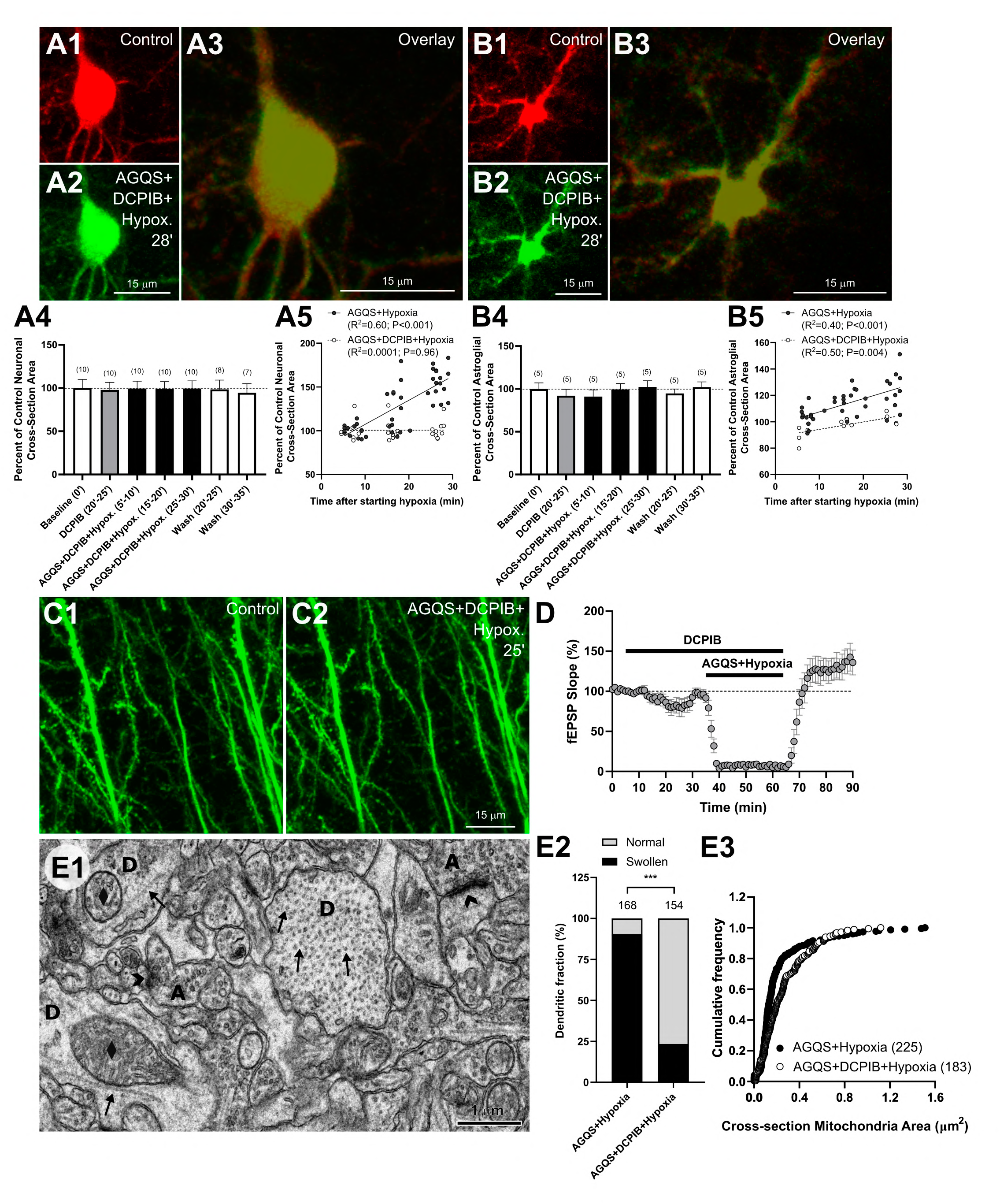
VRAC blockade prevents AGQS-induced neuronal damage during hypoxia. (**A1**-**A3**) A representative color-coded overlay of a CA1 pyramidal neuron shows no changes in neuronal soma size after hypoxia with AGQS and DCPIB (20 µM). (**A4**) Summary from 10 neurons reveals that the size of neuronal somata is unaffected by AGQS during hypoxia if DCPIB is present (F_(8,48)_=0.18, P=0.99, one-way RM ANOVA). (**A5**) Blocking VRAC with DCPIB prevents neuronal swelling elicited by AGQS during hypoxia (F_(1,70)_=5.68, P<0.05, separate-slopes model controlling for the effect of time after start of hypoxia). (**B1**-**B3**) Paired MIP images showing an astrocyte in control (red) and after 28 min of hypoxia with AGQS and DCPIB. Images are overplayed to illustrate the lack of volume change under this experimental condition. (**B4**) Astroglial volume remains stable during hypoxia with AGQS and DCPIB (F_(8,32)_=2.62, P<0.05, one-way RM ANOVA with Dunnett’s post hoc tests that were not significant). (**B5**) Regression line shows a positive correlation between astroglial somata size and duration of hypoxia with AGQS and DCPIB (n=5 astrocytes). Astroglial soma volume during hypoxia with AGQS was significantly reduced in the presence of DCPIB (F_(1,50)_=33.62, P<0.0001, one-way ANCOVA controlling for the effect of time after start of hypoxia). (**C1**, **C2**) Representative MIP images show structurally intact dendrites of CA1 pyramidal neurons in control and hypoxia with DCPIB and AGQS. (**D**) fEPSP was transiently lost during hypoxia with AGQS and DCPIB but fully recovered upon washout and reoxygenation; n=8 slices. (**E1**) Morphologically healthy dendrites (D) with intact cytoplasm and microtubules (arrows) were present in the neuropil of area CA1 of the hippocampus in slices exposed to hypoxia in the presence of AGQS and DCPIB. Synapses (chevrons) were well preserved, and axonal boutons (A) contained synaptic vesicles, but many dendritic mitochondria were swollen (diamonds). (**E2**) Analyses of EM images revealed a significant reduction in a fraction of dendrites with a sign of swelling with DCPIB applied together with AGQS during hypoxia (χ^2_(10)_^=40.54, ***P<0.001, χ^2^ test). (**E3**) Cross-section mitochondrial area was larger during hypoxia with AGQS and DCPIB than without DCPIB (P<0.0001, K-S test).

Importantly, VRAC blockade preserved dendritic morphology (Fig. 6C1, C2), and dendrites remained unbeaded throughout hypoxia with AGQS and DCPIB and during washout with reoxygenation (F_(8, 16)_=1.0, P=0.47, one-way RM ANOVA; data not shown). This lack of dendritic damage was also reflected in the electrophysiological recordings that demonstrated complete functional recovery of the synaptic transmission to the baseline level during washout and reoxygenation (Fig. 6D; P=0.11, Wilcoxon signed-rank test). EM analyses revealed ultrastructurally well-preserved neuropil with intact synapses and axonal boutons (Fig. 6E1). The proportion of swollen dendritic profiles was significantly reduced by about 67% compared to the neuropil exposed to AGQS during hypoxia without a VRAC blocker (Fig. 6E2; P<0.001, χ^2^ test). Surprisingly, the cross-section mitochondrial area was larger during hypoxia and AGQS in the presence of DCPIB than without DCPIB (P<0.0001, K-S test).

These results show that VRAC blockade with DCPIB protects neuropil from AGQS-induced cellular damage during hypoxia and aids in the functional recovery of synaptic transmission.

### GPNA, an ASCT2 blocker, prevents AGQS-induced neuronal and astroglial soma swelling and dendritic damage during hypoxia

The flux of AA through the plasma membrane is tightly regulated by different families of membrane transporters (Bröer and Bröer, 2017). Amino acids contained in the AGQS mixture, except for glycine, are substrates for ASCT2 antiporter that mediates their transport. We hypothesized that inhibition of ASCT2 should reduce the intracellular accumulation of these AA, particularly in astrocytes, diminishing their swelling and lessening neuronal injury during hypoxia with AGQS.

Slices were preincubated for 20 min with GPNA (3 mM) and then exposed to AGQS and hypoxia with GPNA present. Remarkably, neuronal somata swelling was eliminated (Fig. 7A1-A4; P=0.25, one-way RM ANOVA). The absence of swelling was especially evident by the lack of correlation between neuronal somata size and duration of hypoxia with AGQS and GPNA (Fig. 5A5; n=8 neurons, P=0.27, Pearson correlation) and by a significant difference in the slopes of regression lines for AGQS with GPNA and without (P<0.0001, one-way ANCOVA). Likewise, astroglial swelling was also prevented by GPNA treatment (Fig. 7B1-B4; P=0.10, one-way RM ANOVA). Linear regression analysis did not find any correlation between astroglial cross-section area and time passed after starting AGQS and hypoxia with GPNA (Fig. 7B5; n=8 astrocytes, P=0.53, Pearson correlation). The slopes of regression lines for AGQS with GPNA and without differed significantly (Fig. 7B5; P=0.02, one-way ANCOVA), indicating that the inhibition of ASCT2 was sufficient to impede continuous astrocytic swelling induced by AGQS during hypoxia. Dendrites in 6 fields from different mice remained structurally intact in the presence of AGQS with GPNA (Fig. 7C1, 7C2). The lack of dendritic injury has translated to the full recovery of fEPSPs after hypoxia with AGQS and GPNA (Fig. 7D). Actually, fEPSPs were slightly potentiated by 32.7±11% of baseline values after 25 min of washout and reoxygenation (t_(5)_ =2.82, P=0.04, paired t-test).

**Figure 7.**
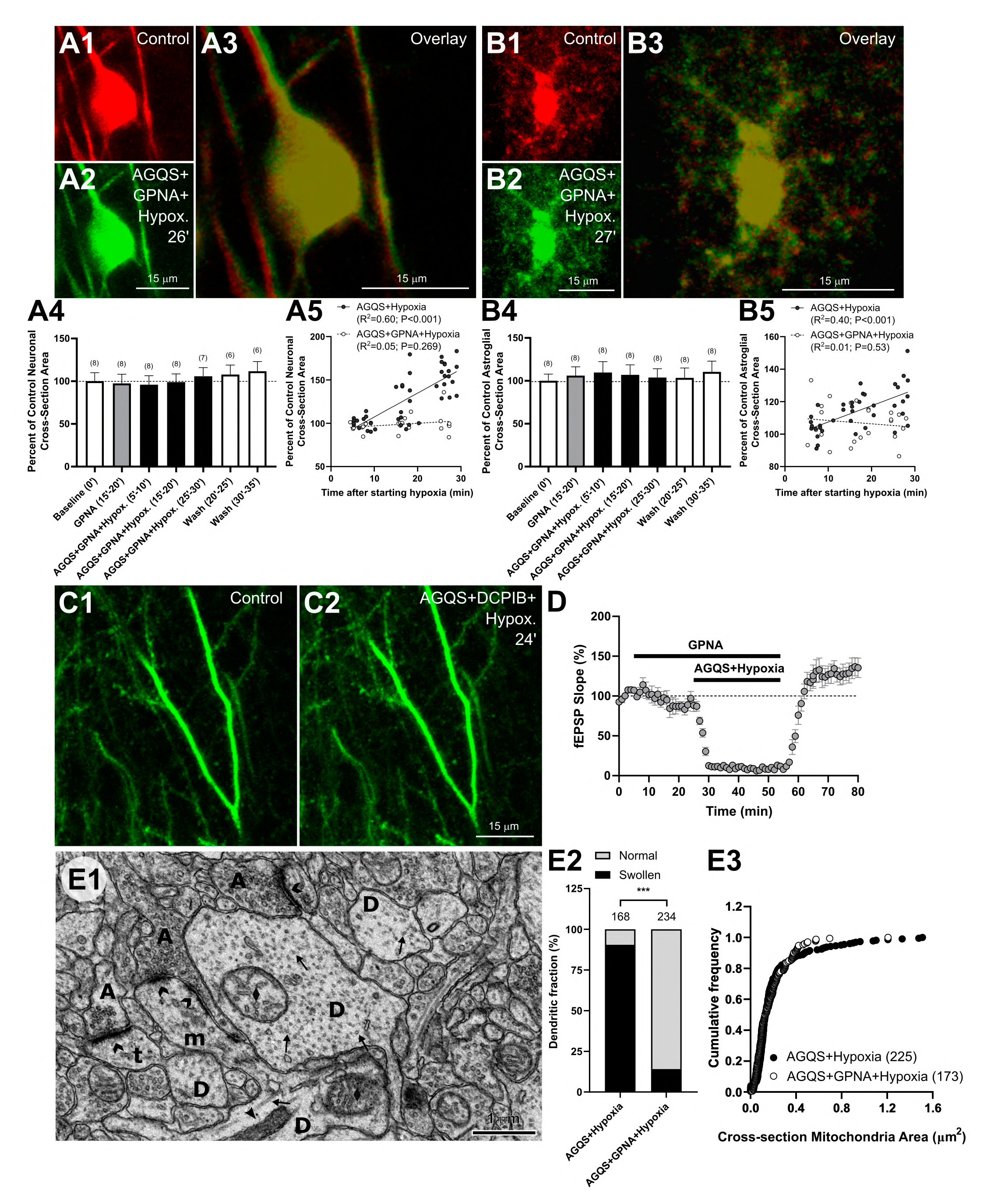
The activity of neutral amino acid transporter ASCT2 is implicated in the onset of damage caused by AGQS in hypoxia. (**A1**-**A3**) Overlay of pyramidal neuronal soma in control and during hypoxia with AGQS and GPNA (3 mM) shows no changes in neuronal cell body volume. (**A4**) Quantification of the size of the neuronal somata reveals no changes during hypoxia with AGQS and GPNA (F_(8,48)_=1.33, P=0.25, one-way RM ANOVA). (**A5**) Treatment with GPNA prevented neuronal swelling elicited during hypoxia in the presence of AGQS (F_(1,64)_=29.55, P<0.0001, one way ANCOVA controlling for the effect of time after start of hypoxia). (**B1**-**B3**) Overlay of astrocyte in control and after 27 min of hypoxia with AGQS and GPNA displays no changes in the size of the astrocyte soma. (**B4**) Astroglial somata size is stable during the hypoxic period in the presence of AGQS and GPNA (F_(8,56)_=1.78, P=0.10, one-way RM ANOVA). (**B5**) Inhibition of ASCT2 transporter with GPNA prevented astroglial swelling during hypoxia with AGQS (F_(1,59)_=5.84, P<0.05, one way ANCOVA controlling for the effect of time after start of hypoxia). (**C1**, **C2**) Dendrites remained healthy in the presence of GPNA during hypoxia with AGQS. (**D**) Synaptic responses were transiently lost in 6 slices exposed to hypoxia in the presence of AGQS and GPNA. Responses recovered and became potentiated after washout and reoxygenation (P=0.04 relative to the baseline, paired t-test). (**E1**) Representative EM image illustrating that GPNA prevented AGQS-induced neuronal injury after 30 min of hypoxia. Dendrites (D) with arrays of microtubules (arrows) were not swollen and contained intact cytoplasm. Some dendritic mitochondria were intact (arrowheads), but some were swollen (diamonds). An example of the well-preserved longitudinally sectioned thin spine (t) is clearly visible. Synapses with morphologically healthy postsynaptic densities (chevron) and axonal boutons (A) full of synaptic vesicles were also evident. (**E2**) EM analyses revealed a significantly smaller fraction of dendrites with signs of swelling when GPNA was added during hypoxia with AGQS (***P<0.001, χ^2^ test). (**E3**) There was no difference in the size of mitochondria after 30 min of hypoxia with AGQS with or without GPNA (P=0.18, K-S test).

In agreement with electrophysiological data and 2PLSM images, electron micrographs revealed largely intact neuropil after 30 min of hypoxia with AGQS and GPNA (Fig. 7E1). The fraction of dendrites with signs of swelling was reduced to ∼14% with GPNA treatment which corresponded to a 76% reduction from hypoxia with AGQS (Fig. 7E2; χ^2_(1)_^ =81.66, P<0.001, χ^2^ test), and it was similar to the proportion of swollen dendrites in neuropil exposed only to hypoxia (χ^2_(1)_^=0.0004, P=0.98, χ^2^ test). Yet, the dendritic mitochondria remained largely swollen even in the presence of GPNA, with no difference in their size after hypoxia and AGQS with or without GPNA (P=0.18, K-S test).

These results confirm that inhibition of ASCT2 protects neuropil from damage induced by AGQS during hypoxia.

A brief summary of our findings is presented in Figure 8.

**Figure 8.**
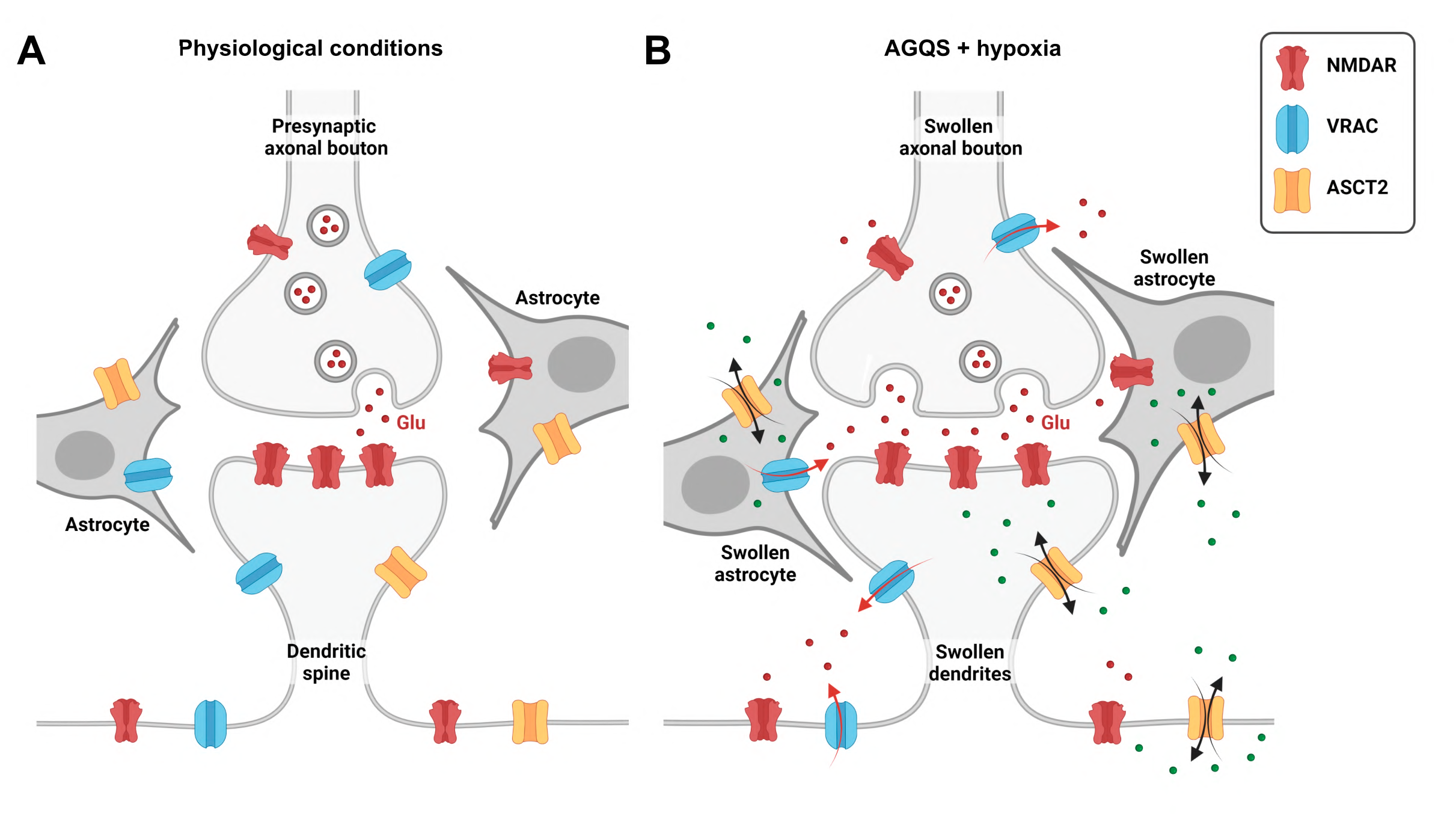
Summary of the hypoxic deleterious effects enhanced by non-excitatory amino acids. (**A**) Astrocytes are tightly coupled with neurons, combining with axons and dendrites to form the tripartite synapse and provide the route for fast glutamate (Glu) clearance under physiological conditions. (**B**) In the presence of AGQS, astrocytes swell due to AGQS accumulation through amino acid transporters, such as the exchanger ASCT2, which is highly expressed in astrocytes. An increase in astroglial volume activates VRAC resulting in the release of gliotransmitters, including glutamate. During hypoxia, glutamate released through VRAC combines with glutamate released by reversed operation of neuronal glutamate transporters and vesicular release and triggers irreversible neuronal damage through an NMDAR-dependent mechanism. Blockade of VRAC or NMDAR or inhibition of ASCT2 prevented NMDAR-dependent neuronal damage and preserved synaptic circuitry. Figure made in BioRender.com.

## Discussion

We used real-time imaging and EM to directly examine structural changes in neurons and astrocytes after exposure to a mixture of four non-excitatory AA (AGQS) at their plasma concentration during normoxic and hypoxic conditions. Our findings are the first to show that during hypoxia, AGQS inflicted irreversible neuronal and astroglial swelling and enduring dendritic damage accompanied by a permanent loss of synaptic transmission.

Synaptic transmission is interrupted in the hypoxic cortex, but recovery of oxygen supply restores synaptic activity without brain damage (Somjen, 2004). Hypoxic failure of synaptic function is mainly due to the activation of presynaptic A1 adenosine receptors that inhibit presynaptic calcium channels and therefore diminish glutamate release (Gundlfinger et al., 2007) exerting protection and allowing recovery of synaptic transmission after hypoxia (Arrigoni et al., 2005). Accordingly, we observed recovery of synaptic potentials with reoxygenation after 30 minutes of hypoxia. However, in the presence of AGQS, the loss of fEPSPs became irreversible.

Our electrophysiological data were supported by 2PLSM and EM findings. Thirty minutes’ hypoxia did not induce astroglial or neuronal soma swelling, and dendrites remained intact. EM confirmed that neuropil was essentially unaffected with well-preserved synapses. There was a slight increase in proportion of dendrites with disrupted microtubules and electron-lucent cytoplasm, indicating some swelling. As expected in hypoxia, mitochondria were swollen (Kovalenko et al., 2006). It is conceivable that many swollen mitochondria and some swollen dendrites could explain the rise in LT during hypoxia (Jarvis et al., 1999). The increase in LT was larger when AGQS were added during hypoxia, signifying cellular swelling. Accordingly, 2PLSM showed increased astroglial and neuronal volume alongside dendritic beading. EM revealed widespread dendritic and mitochondrial enlargement and the presence of disrupted synapses with severely swollen axonal boutons containing clumped synaptic vesicles. No doubt that extensive damage to synaptic circuitry caused a permanent loss of synaptic transmission during hypoxia with AGQS.

In the metabolically compromised cortex, such irreversible injury to synaptic circuitry with astroglial and neuronal swelling and dendritic beading is often caused by SD (Murphy et al., 2008; Risher et al., 2012). Even though sole hypoxia induces SD in slices maintained in an interface chamber (Müller and Somjen, 1999), hypoxia alone cannot trigger SD in submerged slices (Croning and Haddad, 1998). Indeed, SD was never detected in any of our submerged slices either under hypoxia alone or with AGQS, indicating that irreversible cellular swelling and synaptic injury was not resulting from SD but involved a different mechanism enabled by AGQS during hypoxia.

Intracellular accumulation of AA can induce cell swelling as osmotic water accompanies AA. Astrocytes swell due to the expression of aquaporins and water movement through ionic cotransporters (MacAulay, 2021). Astrocytes were swollen in slices exposed to AGQS during normoxia and even more swollen during hypoxia with AGQS, suggesting AA accumulation. Uptake of AGQS could increase intracellular [Na^+^] due to the activity of Na^+^-dependent AA transporters (Bröer and Bröer, 2017). Glycolysis is greatly increased in astrocytes in hypoxia to meet the Na+/K+-ATPase energy demands (Rothman et al., 2022). Yet, it is conceivable that astrocytes were larger during hypoxia as energy deprivation slowed down Na^+^/K^+^-ATPase, leading to the buildup of intracellular Na^+^ accompanied by osmotic water. In contrast, pyramidal neurons lack aquaporins (Papadopoulos and Verkman, 2013) and, therefore, are osmotically tight (Andrew et al., 2007). Thus, osmotically obligated water entry during neuronal AA accumulation is unlikely, explaining the absence of neuronal swelling during normoxia with AGQS. Yet, strongly depolarized neurons swell during seizures (Somjen, 2004), SD (Steffensen et al., 2015), or cooling (Volgushev et al., 2000; Kirov et al., 2004). SD was not implicated in AGQS-induced neuronal swelling during hypoxia, but neurons swelled through an NMDAR-dependent mechanism and thus involved depolarization.

Previously we showed that inhibition of NMDAR rescued the irreversible loss of synaptic potentials induced by a combination of seven non-excitatory AA (AGHQSTU) during hypoxia (Álvarez-Merz et al., 2021). Here, we demonstrated that inhibition of NMDAR during hypoxia with AGQS stopped neuronal soma swelling and dendritic beading without preventing the increase in astroglial soma volume. Synaptic ultrastructure and dendritic integrity were preserved, ensuring recovery of synaptic function. Since most excitatory synapses in the adult brain occur on dendritic spines (Harris and Kater, 1994), dendrites are considered initiation sites of excitotoxic injury leading to neuronal damage and death (Bindokas and Miller, 1995). Indeed, sustained high Ca^2+^ levels coupled with NMDAR activation develop in distal dendrites upon excitotoxic insult and then slowly spread to the soma, triggering acute neuronal injury (Connor et al., 1988; Shuttleworth and Connor, 2001). Dendritic beading is an unmistakable sign of neuronal cytotoxic edema (Kirov et al., 2020), a hallmark of acute neuronal injury (Hori and Carpenter, 1994), and an early sign of cell death pathway activation (Enright et al., 2007). Certainly, intense NMDAR stimulation induces dendritic beading and neuronal soma swelling (Hoskison et al., 2007; Inoue and Okada, 2007). However, when hypoxic slices were exposed only to AGS, no neuronal damage was observed, indicating that the activation of NMDAR was not solely due to the presence of three NMDAR co-agonists: L-alanine, glycine, and L-serine (Pace et al., 1992) in the AGQS mixture. Likewise, L-glutamine can lead to glutamate synthesis in neurons and astrocytes (Bak et al., 2006). Still, L-glutamine alone was insufficient to induce neurotoxicity during hypoxia, indicating that all four AA are necessary.

Normoxic slices accumulate AA after 40 min exposure to AGHQSTU mixture as measured with high-performance liquid chromatography (HPLC) (Álvarez-Merz et al., 2021). It is plausible that during hypoxia, intracellular accumulation of AA with Na^+^ and lactate, generated by glycolysis, underlies substantial astroglial swelling and causes VRAC opening and release of L-glutamate, L-aspartate, and D-serine (Rosenberg et al., 2010; Hyzinski-García et al., 2014). Additionally, glutamate converted from glutamine of the AGQS mixture by intracellular glutaminase could also be released by VRAC, augmenting neurotoxicity. It is noteworthy that VRAC activity is maintained in hypoxia in the presence of glucose, which sustains glycolytic ATP production (Wilson et al., 2019). Once neurotoxicity is established, neurons swell, and neuronal glutamate could also efflux through VRAC (Mongin, 2016). During energy deprivation, neuronal glutamate transporter reversal could also release glutamate (Rossi et al., 2000) adding to the vicious cycle of sustained NMDAR activation and neurotoxicity. Furthermore, in metabolically compromised depolarized neurons, VRAC could mediate the influx of Cl^-^, aggravating swelling and leading to irreversible damage (Inoue and Okada, 2007; Akita and Okada, 2014). In our experiments, the VRAC blocker DCPIB (Zhang et al., 2008) prevented neuronal swelling and dendritic beading produced by AGQS during hypoxia, probably due to the impaired release of excitotoxins through VRAC. Yet, since DCPIB also inhibits glutamate release via connexin hemichannels (Bowens et al., 2013), it cannot be ruled out that this inhibitor was affecting several targets. Surprisingly, inhibition of VRAC reduced astroglial swelling during hypoxia with AGQS. It is plausible that other volume-regulatory mechanisms could compensate for VRAC activity (Wilson and Mongin, 2018). It is also noteworthy that the reduction of the extracellular space concentrates ions and AA, including excitotoxins, and thus, might aggravate hypoxic cell damage. Importantly, the synaptic circuits’ integrity was preserved in the presence of DCPIB, and synaptic function recovered with reoxygenation.

The Na^+^-dependent exchanger ASCT2 (SLC1A5) is expressed in neurons and astrocytes (Bröer et al., 1999; Gliddon et al., 2009) and transports neutral AA, including L-alanine, L-glutamine, and L-serine (Utsunomiya-Tate et al., 1996). ASCT2 plays a major role in the glutamate/GABA-glutamine cycle as one of the main glutamine transporters in the CNS (Scalise et al., 2018). Remarkably, GPNA, an ASCT2 blocker (Foster et al., 2017), was effective in protecting synaptic function while preventing neuronal and astroglial swelling and preserving synaptic ultrastructure during hypoxia with AGQS. ASCT2 mediates both glutamine uptake and efflux in exchange for neutral AA, including L-alanine and L-serine. ASCT2 operates synergistically with other neutral AA transporters from the SLC38 family, such as systems A and N that also transport glutamine, alanine, and serine (Scalise et al., 2018). Perhaps these AA could enter via systems A and N and then alanine and serine efflux via ASCT2 in exchange for glutamine. System A and N are mainly expressed in neurons and astrocytes, respectively (Bröer, 2014; Rubio-Aliaga and Wagner, 2016). However, inhibition of system A in normoxic slices didn’t prevent cellular swelling and tissue accumulation of AA as assessed with electrophysiology and HPLC (Álvarez-Merz et al., 2021). Future studies are necessary to evaluate the involvement of system N. Although GPNA is frequently used as an ASCT2 inhibitor, it is not entirely specific and could also affect systems A and N (Bröer et al., 2016), complicating data interpretation. Conceivably, since AGQS are exchanger substrates, other exchangers, such as Asc-1 (SLC7A10) expressed in neurons and astrocytes (Helboe et al., 2003; Ehmsen et al., 2016), could contribute to neurotoxicity by releasing D-serine (Maucler et al., 2013). Future experiments are warranted to identify AA transporters involved in AGQS-induced damage during hypoxia.

In conclusion, we present a novel mechanism of neuronal injury enacted by non-excitatory AA during hypoxia. Secondary brain damage such as HPC occurs over many days after primary injury. We anticipate that novel therapeutic approaches targeting non-excitatory AA will soon emerge and benefit from our experimental insights into their harmful effects during hypoxic conditions.

## Acknowledgments

The authors thank Libby Perry and Brendan Marshall (Electron Microscopy Core at the Medical College of Georgia) for their assistance with electron microscopy. We thank Dr. Gerald A. Dienel, University of Arkansas for Medical Sciences, Little Rock, Arkansas, for his valuable advice and comments. This work was supported by the National Institutes of Health Grant NS083858 (S.A.K.) and the Ministerio de Universidades, Gobierno de España Grant FPU16/06368 (I.A.M).

## Author contributions

I.A.M, J.M.S., and S.A.K. designed experiments; J.S. provided expert technical support; J.S. and J.M.H.G. provided conceptual advice; I.A.M. performed experiments and analyzed 2PLSM, LT and electrophysiology data with input from S.A.K.; I.V.F. conducted EM analyses; I.A.M. and S.A.K. wrote the manuscript, and all authors edited and approved the manuscript before submission.

## References

Adatia K, Newcombe VFJ, Menon DK (2021) Contusion Progression Following Traumatic Brain Injury: A Review of Clinical and Radiological Predictors, and Influence on Outcome. Neurocrit Care 34:312–324.

Akita T, Okada Y (2014) Characteristics and roles of the volume-sensitive outwardly rectifying (VSOR) anion channel in the central nervous system. Neuroscience 275:211–231.

Alexander JT, Nastuk WL. (1975) Dipping cone to correct optical distortions at liquid surfaces. J Appl Physiol 39:1041–1042.

Álvarez-Merz I, Luengo JG, Muñoz MD, Hernández-Guijo JM, Solís JM. (2021) Hypoxia-induced depression of synaptic transmission becomes irreversible by intracellular accumulation of non-excitatory amino acids. Neuropharmacology 190:108557.

Andrew RD, Labron MW, Boehnke SE, Carnduff L, Kirov SA. (2007) Physiological evidence that pyramidal neurons lack functional water channels. Cereb Cortex 17:787–802.

Andrew RD, Hartings JA, Ayata C, Brennan KC, Dawson-Scully KD, Farkas E, Herreras O, Kirov SA, Müller M, Ollen-Bittle N, Reiffurth C, Revah O, Robertson RM, Shuttleworth CW, Ullah G, Dreier JP. (2022) The Critical Role of Spreading Depolarizations in Early Brain Injury: Consensus and Contention. Neurocrit Care.

Arrigoni E, Crocker AJ, Saper CB, Greene RW, Scammell TE. (2005) Deletion of presynaptic adenosine A1 receptors impairs the recovery of synaptic transmission after hypoxia. Neuroscience 132:575–580.

Bak LK, Schousboe A, Waagepetersen HS. (2006) The glutamate/GABA-glutamine cycle: aspects of transport, neurotransmitter homeostasis and ammonia transfer. J Neurochem 98:641–653.

Bindokas VP, Miller RJ. (1995) Excitotoxic degeneration is initiated at non-random sites in cultured rat cerebellar neurons. J Neurosci 15:6999–7011.

Bowens NH, Dohare P, Kuo YH, Mongin AA. (2013) DCPIB, the proposed selective blocker of volume-regulated anion channels, inhibits several glutamate transport pathways in glial cells. Mol Pharmacol 83:22–32.

Bröer A, Rahimi F, Bröer S. (2016) Deletion of Amino Acid Transporter ASCT2 (SLC1A5) Reveals an Essential Role for Transporters SNAT1 (SLC38A1) and SNAT2 (SLC38A2) to Sustain Glutaminolysis in Cancer Cells. J Biol Chem 291:13194–13205.

Bröer A, Brookes N, Ganapathy V, Dimmer KS, Wagner CA, Lang F, Bröer S. (1999) The astroglial ASCT2 amino acid transporter as a mediator of glutamine efflux. J Neurochem 73:2184–2194.

Bröer S. (2014) The SLC38 family of sodium-amino acid co-transporters. Pflugers Arch 466:155–172.

Bröer S, Bröer A. (2017) Amino acid homeostasis and signalling in mammalian cells and organisms. Biochem J 474:1935–1963.

Chebabo SR, Hester MA, Jing J, Aitken PG, Somjen GG. (1995) Interstitial space, electrical resistance and ion concentrations during hypotonia of rat hippocampal slices. J Physiol 487 (Pt 3):685–697.

Choi DW, Rothman SM. (1990) The role of glutamate neurotoxicity in hypoxic-ischemic neuronal death. Annu Rev Neurosci 13:171–182.

Choi DW, Koh JY, Peters S. (1988) Pharmacology of glutamate neurotoxicity in cortical cell culture: attenuation by NMDA antagonists. J Neurosci 8:185–196.

Connor JA, Wadman WJ, Hockberger PE, Wong RK. (1988) Sustained dendritic gradients of Ca2+ induced by excitatory amino acids in CA1 hippocampal neurons. Science 240:649–653.

Croning MD, Haddad GG. (1998) Comparison of brain slice chamber designs for investigations of oxygen deprivation in vitro. J Neurosci Methods 81:103–111.

Davies ML, Kirov SA, Andrew RD. (2007) Whole isolated neocortical and hippocampal preparations and their use in imaging studies. J Neurosci Methods 166:203–216.

Douglas HA, Callaway JK, Sword J, Kirov SA, Andrew RD. (2011) Potent inhibition of anoxic depolarization by the sodium channel blocker dibucaine. J Neurophysiol 105:1482–1494.

Ehmsen JT, Liu Y, Wang Y, Paladugu N, Johnson AE, Rothstein JD, du Lac S, Mattson MP, Höke A. (2016) The astrocytic transporter SLC7A10 (Asc-1) mediates glycinergic inhibition of spinal cord motor neurons. Sci Rep 6:35592.

Enright LE, Zhang S, Murphy TH. (2007) Fine mapping of the spatial relationship between acute ischemia and dendritic structure indicates selective vulnerability of layer V neuron dendritic tufts within single neurons in vivo. J Cereb Blood Flow Metab 27:1185–1200.

Feng G, Mellor RH, Bernstein M, Keller-Peck C, Nguyen QT, Wallace M, Nerbonne JM, Lichtman JW, Sanes JR. (2000) Imaging neuronal subsets in transgenic mice expressing multiple spectral variants of GFP. Neuron 28:41–51.

Fiala JC. (2005) Reconstruct: a free editor for serial section microscopy. J Microsc 218:52–61.

Foster AC, Rangel-Diaz N, Staubli U, Yang JY, Penjwini M, Viswanath V, Li YX. (2017) Phenylglycine analogs are inhibitors of the neutral amino acid transporters ASCT1 and ASCT2 and enhance NMDA receptor-mediated LTP in rat visual cortex slices. Neuropharmacology 126:70–83.

Gliddon CM, Shao Z, LeMaistre JL, Anderson CM. (2009) Cellular distribution of the neutral amino acid transporter subtype ASCT2 in mouse brain. J Neurochem 108:372–383.

Gundlfinger A, Bischofberger J, Johenning FW, Torvinen M, Schmitz D, Breustedt J. (2007) Adenosine modulates transmission at the hippocampal mossy fibre synapse via direct inhibition of presynaptic calcium channels. J Physiol 582:263–277.

Harris KM, Kater SB. (1994) Dendritic spines: cellular specializations imparting both stability and flexibility to synaptic function. Annu Rev Neurosci 17:341–371.

Hartings JA et al. (2017) The continuum of spreading depolarizations in acute cortical lesion development: Examining Leão’s legacy. J Cereb Blood Flow Metab 37:1571–1594.

Helboe L, Egebjerg J, Møller M, Thomsen C. (2003) Distribution and pharmacology of alanine-serine-cysteine transporter 1 (asc-1) in rodent brain. Eur J Neurosci 18:2227–2238.

Hori N, Carpenter DO. (1994) Functional and morphological changes induced by transient in vivo ischemia. Exp Neurol 129:279–289.

Hoskison MM, Yanagawa Y, Obata K, Shuttleworth CW. (2007) Calcium-dependent NMDA-induced dendritic injury and MAP2 loss in acute hippocampal slices. Neuroscience 145:66–79.

Hossmann KA. (1994) Glutamate-mediated injury in focal cerebral ischemia: the excitotoxin hypothesis revised. Brain Pathol 4:23–36.

Hsu M, Buzsáki G. (1993) Vulnerability of mossy fiber targets in the rat hippocampus to forebrain ischemia. J Neurosci 13:3964–3979.

Hyzinski-García MC, Rudkouskaya A, Mongin AA. (2014) LRRC8A protein is indispensable for swelling-activated and ATP-induced release of excitatory amino acids in rat astrocytes. J Physiol 592:4855–4862.

Inoue H, Okada Y. (2007) Roles of volume-sensitive chloride channel in excitotoxic neuronal injury. J Neurosci 27:1445–1455.

Jackson PS, Strange K. (1993) Volume-sensitive anion channels mediate swelling-activated inositol and taurine efflux. Am J Physiol 265:C1489–1500.

Jarvis CR, Lilge L, Vipond GJ, Andrew RD. (1999) Interpretation of intrinsic optical signals and calcein fluorescence during acute excitotoxic insult in the hippocampal slice. Neuroimage 10:357–372.

Jensen FE, Harris KM. (1989) Preservation of neuronal ultrastructure in hippocampal slices using rapid microwave-enhanced fixation. J Neurosci Methods 29:217–230.

Kirov SA, Sorra KE, Harris KM. (1999) Slices have more synapses than perfusion-fixed hippocampus from both young and mature rats. J Neurosci 19:2876–2886.

Kirov SA, Fomitcheva IV, Sword J. (2020) Rapid Neuronal Ultrastructure Disruption and Recovery during Spreading Depolarization-Induced Cytotoxic Edema. Cereb Cortex 30:5517–5531.

Kirov SA, Petrak LJ, Fiala JC, Harris KM. (2004) Dendritic spines disappear with chilling but proliferate excessively upon rewarming of mature hippocampus. Neuroscience 127:69–80.

Kovalenko T, Osadchenko I, Nikonenko A, Lushnikova I, Voronin K, Nikonenko I, Muller D, Skibo G. (2006) Ischemia-induced modifications in hippocampal CA1 stratum radiatum excitatory synapses. Hippocampus 16:814–825.

Kurland D, Hong C, Aarabi B, Gerzanich V, Simard JM. (2012) Hemorrhagic progression of a contusion after traumatic brain injury: a review. J Neurotrauma 29:19–31.

Lerma J, Herranz AS, Herreras O, Abraira V, Martín del Río R. (1986) In vivo determination of extracellular concentration of amino acids in the rat hippocampus. A method based on brain dialysis and computerized analysis. Brain Res 384:145–155.

MacAulay N. (2021) Molecular mechanisms of brain water transport. Nat Rev Neurosci 22:326–344.

Maucler C, Pernot P, Vasylieva N, Pollegioni L, Marinesco S. (2013) In vivo D-serine hetero-exchange through alanine-serine-cysteine (ASC) transporters detected by microelectrode biosensors. ACS Chem Neurosci 4:772–781.

Mongin AA. (2016) Volume-regulated anion channel--a frenemy within the brain. Pflugers Arch 468:421–441.

Müller M, Somjen GG. (1999) Intrinsic optical signals in rat hippocampal slices during hypoxia-induced spreading depression-like depolarization. J Neurophysiol 82:1818–1831.

Murphy TH, Li P, Betts K, Liu R. (2008) Two-photon imaging of stroke onset in vivo reveals that NMDA-receptor independent ischemic depolarization is the major cause of rapid reversible damage to dendrites and spines. J Neurosci 28:1756–1772.

Nishimura F, Nishihara M, Mori M, Torii K, Takahashi M. (1995) Excitability of neurons in the ventromedial nucleus in rat hypothalamic slices: modulation by amino acids at cerebrospinal fluid levels. Brain Res 691:217–222.

Nolte C, Matyash M, Pivneva T, Schipke CG, Ohlemeyer C, Hanisch UK, Kirchhoff F, Kettenmann H. (2001) GFAP promoter-controlled EGFP-expressing transgenic mice: a tool to visualize astrocytes and astrogliosis in living brain tissue. Glia 33:72–86.

Pace JR, Martin BM, Paul SM, Rogawski MA. (1992) High concentrations of neutral amino acids activate NMDA receptor currents in rat hippocampal neurons. Neurosci Lett 141:97–100.

Papadopoulos MC, Verkman AS. (2013) Aquaporin water channels in the nervous system. Nat Rev Neurosci 14:265–277.

Risher WC, Andrew RD, Kirov SA. (2009) Real-time passive volume responses of astrocytes to acute osmotic and ischemic stress in cortical slices and in vivo revealed by two-photon microscopy. Glia 57:207–221.

Risher WC, Croom D, Kirov SA. (2012) Persistent astroglial swelling accompanies rapid reversible dendritic injury during stroke-induced spreading depolarizations. Glia 60:1709–1720.

Risher WC, Ard D, Yuan J, Kirov SA. (2010) Recurrent spontaneous spreading depolarizations facilitate acute dendritic injury in the ischemic penumbra. J Neurosci 30:9859–9868.

Risher WC, Lee MR, Fomitcheva IV, Hess DC, Kirov SA. (2011) Dibucaine mitigates spreading depolarization in human neocortical slices and prevents acute dendritic injury in the ischemic rodent neocortex. PLoS One 6:e22351.

Rosen AS, Andrew RD. (1990) Osmotic effects upon excitability in rat neocortical slices. Neuroscience 38:579–590.

Rosenberg D, Kartvelishvily E, Shleper M, Klinker CM, Bowser MT, Wolosker H. (2010) Neuronal release of D-serine: a physiological pathway controlling extracellular D-serine concentration. Faseb j 24:2951–2961.

Rossi DJ, Oshima T, Attwell D. (2000) Glutamate release in severe brain ischaemia is mainly by reversed uptake. Nature 403:316–321.

Rothman DL, Dienel GA, Behar KL, Hyder F, DiNuzzo M, Giove F, Mangia S. (2022) Glucose sparing by glycogenolysis (GSG) determines the relationship between brain metabolism and neurotransmission. J Cereb Blood Flow Metab 42:844–860.

Rubio-Aliaga I, Wagner CA. (2016) Regulation and function of the SLC38A3/SNAT3 glutamine transporter. Channels (Austin) 10:440–452.

Scalise M, Pochini L, Console L, Losso MA, Indiveri C. (2018) The Human SLC1A5 (ASCT2) Amino Acid Transporter: From Function to Structure and Role in Cell Biology. 6.

Schipke CG, Ohlemeyer C, Matyash M, Nolte C, Kettenmann H, Kirchhoff F. (2001) Astrocytes of the mouse neocortex express functional N-methyl-D-aspartate receptors. Faseb j 15:1270–1272.

Shuttleworth CW, Connor JA. (2001) Strain-dependent differences in calcium signaling predict excitotoxicity in murine hippocampal neurons. J Neurosci 21:4225–4236.

Somjen GG. (2004) Ions in the Brain: Normal Function, Seizures, and Stroke: Oxford University Press.

Steffensen AB, Sword J, Croom D, Kirov SA, MacAulay N. (2015) Chloride Cotransporters as a Molecular Mechanism underlying Spreading Depolarization-Induced Dendritic Beading. J Neurosci 35:12172–12187.

Sword J, Croom D, Wang PL, Thompson RJ, Kirov SA. (2017) Neuronal pannexin-1 channels are not molecular routes of water influx during spreading depolarization-induced dendritic beading. J Cereb Blood Flow Metab 37:1626–1633.

Utsunomiya-Tate N, Endou H, Kanai Y. (1996) Cloning and functional characterization of a system ASC-like Na+-dependent neutral amino acid transporter. J Biol Chem 271:14883–14890.

Vander Jagt TA, Connor JA, Shuttleworth CW. (2008) Localized loss of Ca2+ homeostasis in neuronal dendrites is a downstream consequence of metabolic compromise during extended NMDA exposures. J Neurosci 28:5029–5039.

Verkhratsky A, Chvátal A. (2020) NMDA Receptors in Astrocytes. Neurochem Res 45:122–133.

Volgushev M, Vidyasagar TR, Chistiakova M, Yousef T, Eysel UT. (2000) Membrane properties and spike generation in rat visual cortical cells during reversible cooling. J Physiol 522 Pt 1:59–76.

Weiss MD, Derazi S, Kilberg MS, Anderson KJ. (2001) Ontogeny and localization of the neutral amino acid transporter ASCT1 in rat brain. Brain Res Dev Brain Res 130:183–190.

Wilson CS, Mongin AA. (2018) Cell Volume Control in Healthy Brain and Neuropathologies. Curr Top Membr 81:385–455.

Wilson CS, Bach MD, Ashkavand Z, Norman KR, Martino N, Adam AP, Mongin AA. (2019) Metabolic constraints of swelling-activated glutamate release in astrocytes and their implication for ischemic tissue damage. J Neurochem 151:255–272.

Yan J, Bengtson CP, Buchthal B, Hagenston AM, Bading H. (2020) Coupling of NMDA receptors and TRPM4 guides discovery of unconventional neuroprotectants. Science 370.

Zhang Y, Zhang H, Feustel PJ, Kimelberg HK. (2008) DCPIB, a specific inhibitor of volume regulated anion channels (VRACs), reduces infarct size in MCAo and the release of glutamate in the ischemic cortical penumbra. Exp Neurol 210:514–520.

